# An Igh novel enhancer modulates antigen receptor diversity by determining locus conformation

**DOI:** 10.1101/2022.05.23.492988

**Authors:** Khalid H. Bhat, Saurabh Priyardarshi, Sarah Naiyer, X. Qu, Hammad Farooq, Eden Kleiman, J. Xu, X. Lei, Jose F. Cantillo, Robert Wuerffel, Nicole Baumgarth, Jie Liang, Ann J. Feeney, Amy L. Kenter

## Abstract

The Igh locus is organized into a developmentally regulated topologically associated domain (TAD) that is divided into subTADs. Here we identify a series of novel enhancers (NEs) that collaborate to configure the locus, determine transcriptional potential in over a hundred functional V_H_ genes and their usage in V(D)J recombination. NE1 engages in a network of long-range interactions that interconnect the subTADs and the recombination center at the D_H_J_H_ gene cluster. Deletion of NE1 alters discrete chromatin loops, higher order locus conformation, locus-wide V_H_ gene transcription and regional V gene utilization that is linked to a greatly reduced splenic B1 B cell compartment. NE1 blocks long-range loop extrusion that in turn contributes to locus contraction and determines the proximity of distant V_H_ genes to the recombination center. NE1 is a critical architectural element that coordinates chromatin conformational states that favor V_H_ gene transcription or V(D)J rearrangement.

## INTRODUCTION

Progenitor B cells must develop a diverse antibody receptor repertoire to provide protection against a wide range of antigens and pathogens. Each mature B cell has a unique Ig receptor, created via V(D)J recombination. In each pro-B cell, one of the ∼100 functional Igh locus V_H_ genes must recombine with a rearranged DJ_H_ element, which itself is assembled from one of 8-12 D_H_ and one of 4 J_H_ gene segments (Schatz and Ji, 2011). These observations raise the question of how the distal V_H_ gene segments, located up to 2.5 Mb from the DJ_H_ join, engage in rearrangement since V_H_ genes throughout the locus are used in the repertoire. In pro-B cells, the Igh locus contracts to juxtapose distal V_H_ genes next to proximal D_H_ segments to promote V(D)J joining (Fuxa et al., 2004; Jhunjhunwala et al., 2008; Kosak et al., 2002; Roldan et al., 2005; Sayegh et al., 2005). Igh locus contraction requires the lineage commitment factor, Pax5 and the transcriptional regulator, YY1 (Fuxa *et al*., 2004; Liu et al., 2007). However, the mechanistic relationship of locus contraction to looping interactions and DNA motifs specifying the spatial organization of the Igh locus remain largely undefined.

Chromatin is organized at the Mb scale into topological associating domains (TADs) that encompass spatial neighborhoods of high frequency chromatin interactions (Dixon et al., 2012; Hou et al., 2012; Nora et al., 2012; Sexton et al., 2012). TAD organization reflects the functional partition of chromatin regions by transcriptional activity (Dixon *et al*., 2012; Sexton *et al*., 2012), histone modifications (Dixon *et al*., 2012; Nora *et al*., 2012; Rao et al., 2014; Sexton *et al*., 2012), and replication timing (Pope et al., 2014) implying a link between function and genome structure. The Igh locus is contained within a 2.9-Mb TAD that has few compartment interactions in pro-B cells (Montefiori et al., 2016).

TADs are frequently anchored by DNA elements bound by the architectural protein CTCF and its partner the SMC cohesin complex (Rao *et al*., 2014; Rowley and Corces, 2018). The observation that CTCF binding elements (CBEs) situated at TAD boundaries are often in a convergent orientation (Rao *et al*., 2014) has led to the proposition that chromatin loops are formed by an extrusion mechanism mediated by cohesin (Fudenberg et al., 2016; Nasmyth, 2001; Nichols and Corces, 2015; Sanborn et al., 2015). Studies have implicated cohesin mediated loop extrusion in generating Igh locus contraction (Ba et al., 2020; Hill et al., 2020) and as a mechanism for a RAG scanning model implicated in VDJ gene assembly (Dai et al., 2021; Jain et al., 2018; Zhang et al., 2019). However, Igh locus topology has also been described as configured by three large chromatin loops in pre-pro-B cells that become intermingled and provide equal access of the D_H_-distal and -proximal V_H_ gene segments with rearranged 3’ D_H_J_H_ in pro-B cells (Jhunjhunwala *et al*., 2008). Our previous studies of Igh locus chromatin architecture defined chromatin loops and anchor sites that contribute to locus topology (Montefiori *et al*., 2016). How can we reconcile a specific chromatin loop model of Igh locus conformation and the plethora of loops formed by loop extrusion? Loop extrusion also provides an explanation for how enhancer-enhancer (E-E) and promoter-enhancer (Pr-E) anchored loops may form within a TAD (Fudenberg *et al*., 2016; Hsieh et al., 2020; Schwarzer et al., 2017). We favor an integrated view in which loop extrusion sets up a series of chromatin loops anchored at specific regulatory elements in the Igh locus. Our new studies presented here provide evidence for this perspective.

We have developed chromatin conformation and single cell imaging approaches to define essential DNA motifs that anchor chromatin loops and drive the 3D conformation of the Igh locus. We report discovery of four Igh novel enhancers (NE1-4) and demonstrate that NE1 has structural and functional impact on the Igh locus. NE1 participates in an enhancer interactome, determines locus topology, modulates V_H_ gene transcription and regional V_H_ gene usage during VDJ recombination. Our findings reveal that NE1 constrains loop extrusion and thereby determines chromatin conformational states that favor either VDJ recombination or V_H_ gene transcription.

## RESULTS

### Four novel enhancers identified in the Igh locus

The Igh TAD is divided into three sub-TADs linked by Site I of sub-TAD A, Friend of Site Ia (FrOStIa) and FrOStIb of sub-TAD B, and Sites II, II.5 and III all in sub-TAD C (Montefiori *et al*., 2016) (fig. 1A,G). To define motifs that anchor Igh chromatin loops, Site I and FrOStIa were scanned for distinct transcriptional and epigenetic features (Suppl. Table 1). The highly transcribed V_H_14-2 gene promoter (Pr) is prominently decorated with histone 3 lysine 4 methyl 1 (H3K4me1), H3K4me3 and is bound by the co-activator MED1 and TFs Pax5 and IRF4 consistent with its transcriptional activity and lacks CTCF and RAD21 (fig. 1B, left panel). H3K4me3 marks transcriptionally active Prs (Heintzman et al., 2007). NE1 is marked by H3K27Ac and H3K4me1 and bound by TFs PAX5, IRF4, IKAROS, E2A, PU.1, and transcriptional co-activators MED1, BRG1 and p300 (fig. 1C). H3K4me1 is associated with active and poised enhancers while H3K27Ac distinguishes active enhancers (reviewed in (Calo and Wysocka, 2013)). NE1 is flanked by a CTCF binding element (CBE) that is co-bound by CTCF and RAD21 (fig. 1A, C). Detection of NE1 prompted us to screen the Igh locus for other potential NEs. We detected a series of NEs including NE2, NE3, and NE4 (red rectangles) with a TF binding profile similar to NE1 that are interspersed among the intermediate and distal V_H_ gene exons (fig. 1A, C)(Suppl. fig. 1). NE2 is located within FrOStIb in sub-TAD B and NE3, and NE4 are positioned between Site II and Site II.5 in sub-TAD C (fig. 1A,C). We consider the possibility that V_H_14-2 Pr and the NEs play a role in V_H_ gene transcription and/or locus topology.

**Figure 1.**
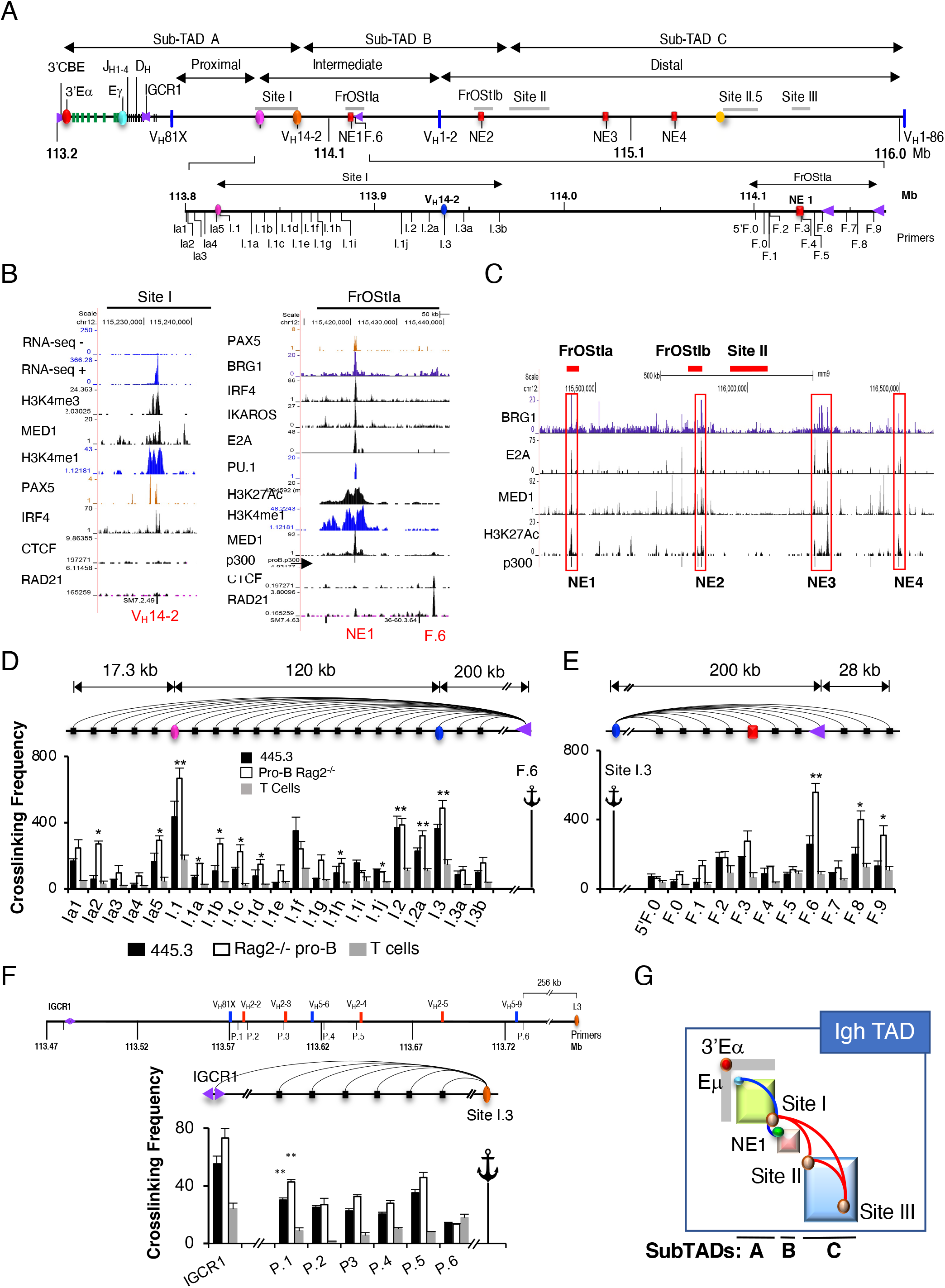
Identification of Site I and FrOStIa loop anchors motifs. All genomic coordinates (chr12, mm10). **A)** Schematic of the Igh locus. **B)** Chip-seq and RNA-seq analyses for the Site I V_H_14-2 gene. **C)** ChIP-seq studies identify NE1, NE2, NE3 and NE4 (red rectangles). **D-F)** 3C assays. Arcs indicate individual 3C assays, primers are identified below the histograms, and anchor probes are identified (anchor symbol). Average crosslinking frequencies are from at least two independent samples and SEMs are shown. Two-tail Student’s t test. **D)** 3C assays analyzing Site I and anchored at F.6. **E)** 3C assays analyzing FrOStIa and anchored at Site I.3. **F)** (*Upper panel*) Schematic of subTAD A showing a subset of V_H_ genes and 3C primers (P). (*Lower panel*) 3C assays using the Site I.3 (I.3) anchor and P.1-P.6 primers. **G)** Schematic of the Igh locus showing sub-TADs and looping interactions (Montefiori *et al*., 2016).

### Identification of Site I and FrOStIa loop anchors

We examined the spatial organization of Site I and FrOStIa in CD19^+^ Rag2^-/-^ pro-B cells, the Abelson transformed (Abl-t) Rag1 deficient pro-B cell line, 445.3 and ConA activated splenic T cells using the 3C anchors, FrOStIa F.6 and NE1 (fig. 1C) (Suppl. fig. 1). The F.6 3C fragment contains CBE and is located ∼15 kb upstream of NE1 (fig. 1B). In both cases, looping interactions with 3C fragments Site I.1, I.2 and I.3 (V_H_14-2 gene) were elevated in pro-B cell specific chromatin templates as compared to that found in T cells (fig. 1D) (Suppl. fig. 1B). Likewise, the Site I.3 anchor probe, which harbors the V_H_14-2 gene, associates with F.2-F.3 (NE1) and F.6 (CBE) and F.8-F.9 in pro-B cell specific fashion (fig. 1E). Thus, the Site I.3 fragment is a major interaction partner with the NE1 and F.6 (CBE). It was difficult to determine candidate loop anchors in Site I.1 and I.2 due to an unremarkable epigenetic landscape and TF binding profile.

To determine whether the Site I.3 fragment also associates with D_H_ proximal V_H_ genes we analyzed locations close to V_H_ segments (P.1-P.6) and IGCR1 in chromatin from Rag2^-/-^ pro-B cells, the 445.3 line, and ConA activated splenic T cells in 3C assays (fig. 1F, upper panel). IGCR1, composed of two CBE in divergent orientation, is a boundary element that separates the D_H_-J_H_ clusters from the most proximal V_H_ genes (Degner et al., 2009; Jain *et al*., 2018; Qiu et al., 2018) (fig. 1A). Site I.3 contacts P.1-P.5 and IGCR1 but not P.6 in pro-B cell specific chromatin and not in T cells (fig. 1F, lower panel). Thus, Site I.3 engages in a multiplicity of interactions spanning sub-TAD A (blue arc) and may act as a bridge between the D_H_ proximal V_H_ genes and NE1/F.6 located in subTAD B (fig. 1G).

### Disruption of a V_H_ promoter, NE and CBE diminishes Igh looping and transcription

To establish a cell culture model to study the involvement of specific transcriptional elements (TE) and CBE motifs in Igh locus function we employed genome editing to generate bi-allelic identical deletions of the V_H_14-2 promoter (Pr), NE1, F.6 CBE, NE2 and Site I.1 in the 445.3.11 sub-line (fig. 2A)(Suppl. fig. 2). V_H_ Prs are composed of a polypyrimidine tract, heptamer and octamer and deletion of the octamer/heptamer is sufficient to abolish Pr activity (Eaton and Calame, 1987). CTCF binding at the F.6 CBE in the control- and Site I.1 CBE KO lines was detected whereas binding was abolished in the F.6 CBE KO lines in ChIP assays (fig. 2B). Because there was little epigenetic guidance to identify potential loop anchor motifs within Site I.1 we arbitrarily constructed deletions centered on the V_H_2-8 exon and an adjacent CBE (Suppl. fig. 2E). Site I.3:F.6 looping interactions were dependent on the integrity of the V_H_14-2 Pr, F.6 CBE and NE1 motifs but not on NE2 or deletions around the V_H_2-8 gene in Site I.1 (fig. 2C) (Suppl. fig. 2F,G). F.6:Site I.1 and Site I.3:Site I.1 contacts required an intact F.6 CBE and V_H_14-2 Pr, respectively, indicating the autonomy of each these interactions (fig. 2C). Thus, the V_H_14-2 Pr, NE1 and F.6 CBE coordinate Site I:FrOStIa looping.

**Figure 2.**
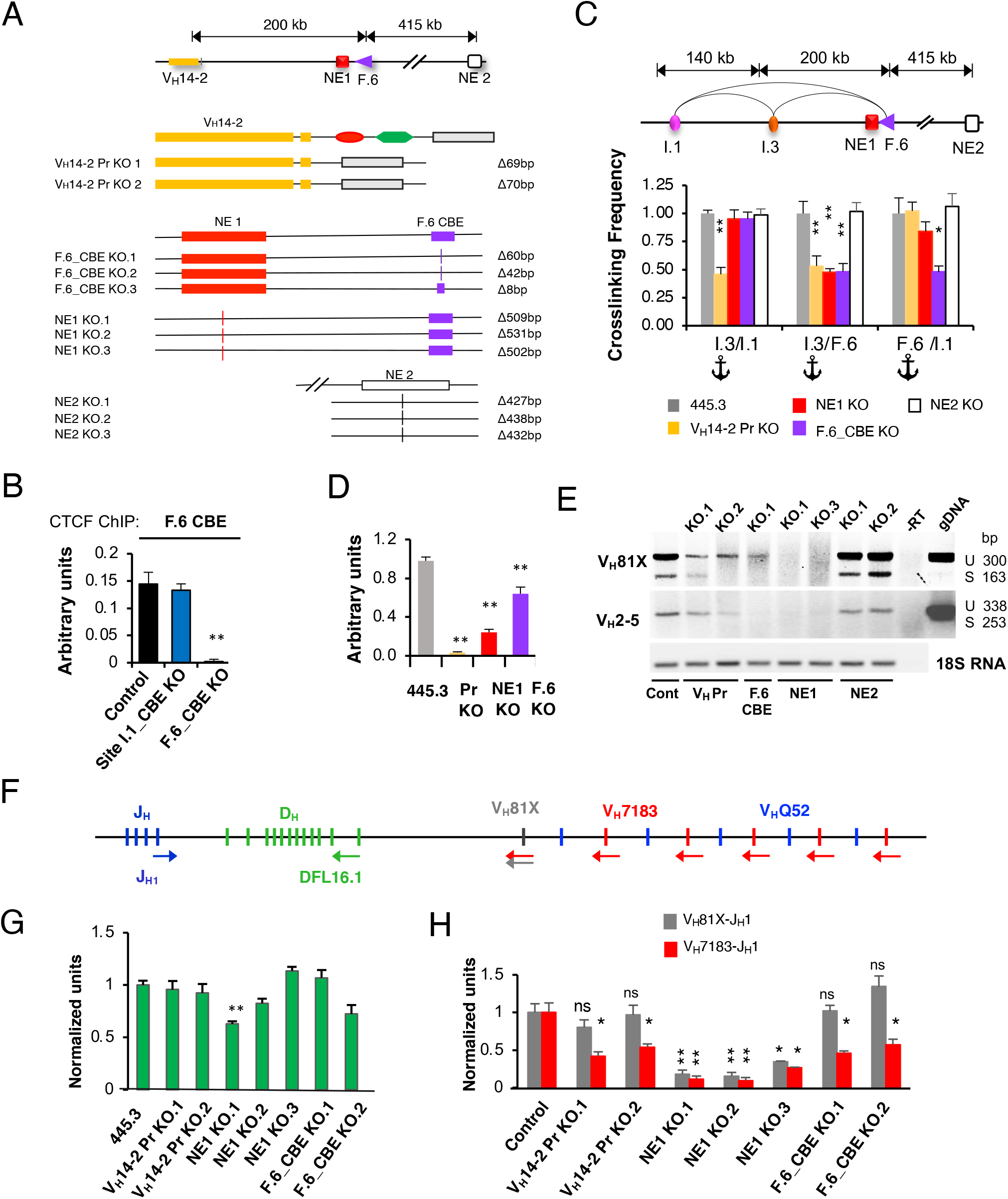
Igh locus function is dependent on a V_H_ promoter, a NE and CBE in Abl-t pro-B cells. P values are from two-tail Student’s t test. **A)** (*Upper panel*) The Igh locus from the V_H_14-2 gene to NE2. (*Lower panel)* Biallelic identical deletions constructed in the Abl-t 445.3.11 pro-B cell line. **B)** ChIP assays for CTCF binding at the F.6 CBE. **C)** (*Upper panel*) Map of genomic distances. (*Lower panel)* Normalized 3C crosslinking frequencies, anchored at Site I.3 or F.6 using chromatin from control and KO lines. **D)** QRT-PCR assays for the V_H_14-2 gene using 18S RNA as a loading control. **E)** RT-PCR of V_H_81X and V_H_2-5 GLTs and the 18S RNA loading control were harvested at 32 and 28 cycles, respectively. Cells were STI-571 (2.5 mM) treated for 48h. **F)** Schematic of the D_H_J_H_ clusters and proximal V_H_ gene segments (vertical bars), with primer positions (colored arrows) shown. **G**,**H)** Cell lines were stably complemented with RAG1 and STI-517 treated. **G)** D-J rearrangements in gDNA were analyzed by qPCR with DFL16.1 and J_H_1 primers. **H)** Normalized qRT-PCR data for V->DJ rearrangement using V_H_81X and V_H_7183 primers in combination with J_H_1.

To validate NE1 enhancer function we tested the transcriptional effects of deletion on V_H_ germline transcript (GLT) expression. V_H_14-2 GLT expression was abolished or significantly diminished in the V_H_14-2 Pr-, NE1- and F.6-KO lines, respectively, as compared to the control demonstrating that the transcription of this GLT is both Pr- and NE1-dependent (fig. 2D). We studied two additional V_H_ genes in 445.3.11 cells that were arrested in G1 phase by treatment with STI571 to induce Ig gene transcription (Muljo and Schlissel, 2003). V_H_81X is a member of the V_H_7183 family and the first functional D_H_ proximal V_H_ gene and V_H_2-5, is a member of the V_H_Q52 family and located in sub-TAD A (fig. 1A). V_H_ GLTs, composed of a 5’ leader sequence and V_H_ exon are often found as unspliced and spliced where the PCR amplification of gDNA reflects the MW of the unspliced GLT (fig. 2E). The V_H_81X (unspliced, 300 bp; spliced, 163 bp) and V_H_2-5 (unspliced, 338 bp) GLTs are clearly present in the 445.3.11 control and in the NE2 KO lines (fig. 2E). However, these transcripts are abolished in NE1 KO cells, and are diminished in V_H_14-2 Pr- or F.6_CBE KO lines indicating that 1) NE1 is essential for transcription of at least several V_H_ genes and 2) that the V_H_14-2 Pr and F.6 CBE elements may contribute to NE1 function, by facilitating its spatial proximity to V_H_ exons in subTAD-A.

### V(D)J recombination requires specific loop anchors in Abl-t pro-B cell lines

To investigate the influence of TE and CBE KOs on V(D)J recombination we studied D->J_H_ and V_H_->DJ_H_ rearrangements in Rag1 deficient Abl-t 445.3.11 control and KO lines. D->J_H_ recombination reflects the activity of the Eμ proximal recombination center (RC) that forms over the J_H_ cluster and to which RAG1/2 is recruited (Ji et al., 2010). Cells were stably complemented with a RAG1 expression construct, viably arrested in G1 phase by treatment with STI-571 to induce endogenous RAG2 and VDJ recombination (Bredemeyer et al., 2006). D->J_H_ rearrangements, assessed in gDNA using the DFL.16 and J_H_1 primers, were largely comparable in control and KO lines indicating that the first step in V(D)J recombination is intact and independent of TE and CBE deletions (fig. 2 F,G). Slightly reduced D->J_H_ recombination in NE1 KO.1 may be attributable to subclone variability (fig. 2 F,G).

V_H_81X, a member of the V_H_7183 family is the most highly used VH gene in Abl-t pro-B cell lines (Lee et al., 2014) and in mouse C57Bl/6 pro-B cells (Bolland et al., 2016) whereas the distal VHJ558->DJ_H_ recombination is rare in Abl-t pro-B cell lines (Lee *et al*., 2014). V_H_7183->DJ_H_ rearrangements were analyzed in qRT-PCR using two forward primer sets specific for V_H_81X-(gray arrow), and a pan specific V_H_7183 primer (red arrow) in combination with the reverse J_H_1 primer (blue arrow) (fig. 2F). Although V_H_7183-rearrangement levels were strikingly reduced in all Abl-t pro-B cell lines harboring TE and CBE deletions, V_H_81X rearrangements were significantly diminished only in NE1 KO lines (fig. 2I). To address this difference, we asked if V_H_ gene expression was correlated with the propensity to engage in rearrangement. V_H_81X transcription was reduced but not abolished in V_H_14-2 Pr- and F.6 CBE KO lines whereas V_H_81X GLTs were undetectable in the NE1 KO lines suggesting that even low-level transcription may contribute to V_H_81X->DJ_H_ recombination (fig. 2E). Thus, loss of NE1 led to the deepest deficits for both transcription and recombination relative to other KOs (fig. 2 E,I). However, reduced transcription may not cause reduced recombination since V_H_14-2 Pr, NE1, and F.6 CBE motifs also mediate long range looping interactions that could determine the proximity of V_H_ genes to the RC as a precursor to V_H_->DJ_H_ recombination. In so doing, transcription may be a byproduct of V_H_ gene segment proximity to Eμ rather than a precondition to recombination. Nevertheless, the ablation of V_H_81X transcription correlates with abolished V_H_81X->DJ recombination in the NE1 KO lines.

### A Eμ-V_H_Pr-NE1 chromatin hub mediates Igh locus architecture

Efficient V_H_->DJ_H_ recombination, transcription and chromatin loop formation all require intact V_H_14-2 Pr, NE1 and F.6 CBE motifs suggesting that these elements organize essential structure/function relationships in the Igh locus. Average spatial distance between markers generally increase as a function of genomic distance in 3D DNA FISH. However, in pro-B cells mean spatial distances between markers arrayed across the Igh locus flattened upon increasing genomic distance indicating compaction of the locus and proximity of the markers (Jhunjhunwala *et al*., 2008). To explore the topology of the Igh locus we measured spatial distances between Eμ and 3’Eα, V_H_14-2 Pr (Site I.3) and NE1 using short FISH probes (4.8-7.5 kb) that are separated by 220 kb (Eμ-3’Eα), 568 kb (Eμ-Site I.3) and 190 kb (Site I.3-NE1), respectively (fig. 3A,B). The mean spatial distances separating the Eμ anchor probe from the 3’Eα and Site I.3 probes increased as a function of genomic distance and then leveled off between Site I.3 and NE1 in controls (fig. 3C). Notably, the mean spatial distance increased in NE1 and F.6_CBE KO lines as compared to the control indicating substantial decompaction of the locus upon loss of these elements (fig. 3C,D). Moreover, the increased mean spatial distances involving Eμ and 3’Eα in the NE1- and F.6_CBE KO lines were confirmed in 3C assays which showed reduced crosslinking frequency for Eμ:3’Eα: as compared to the control line (fig. 3E). Deletion of the V_H_14-2 Pr led to a dramatic increase of average spatial distance between Eμ and NE1 whereas Eμ and Site I.3 were more compacted relative to the control (fig. 3C,D). Thus, Eμ:Site I.3 association is not mediated by direct interaction with the V_H_14-2 Pr and must therefore be supported by another element. We conclude that NE1 and F.6_CBE have a profound influence on locus topology in subTADs A-B and that the V_H_14-2 Pr:NE1 interaction serves to bridge NE1 to Eμ.

**Figure 3.**
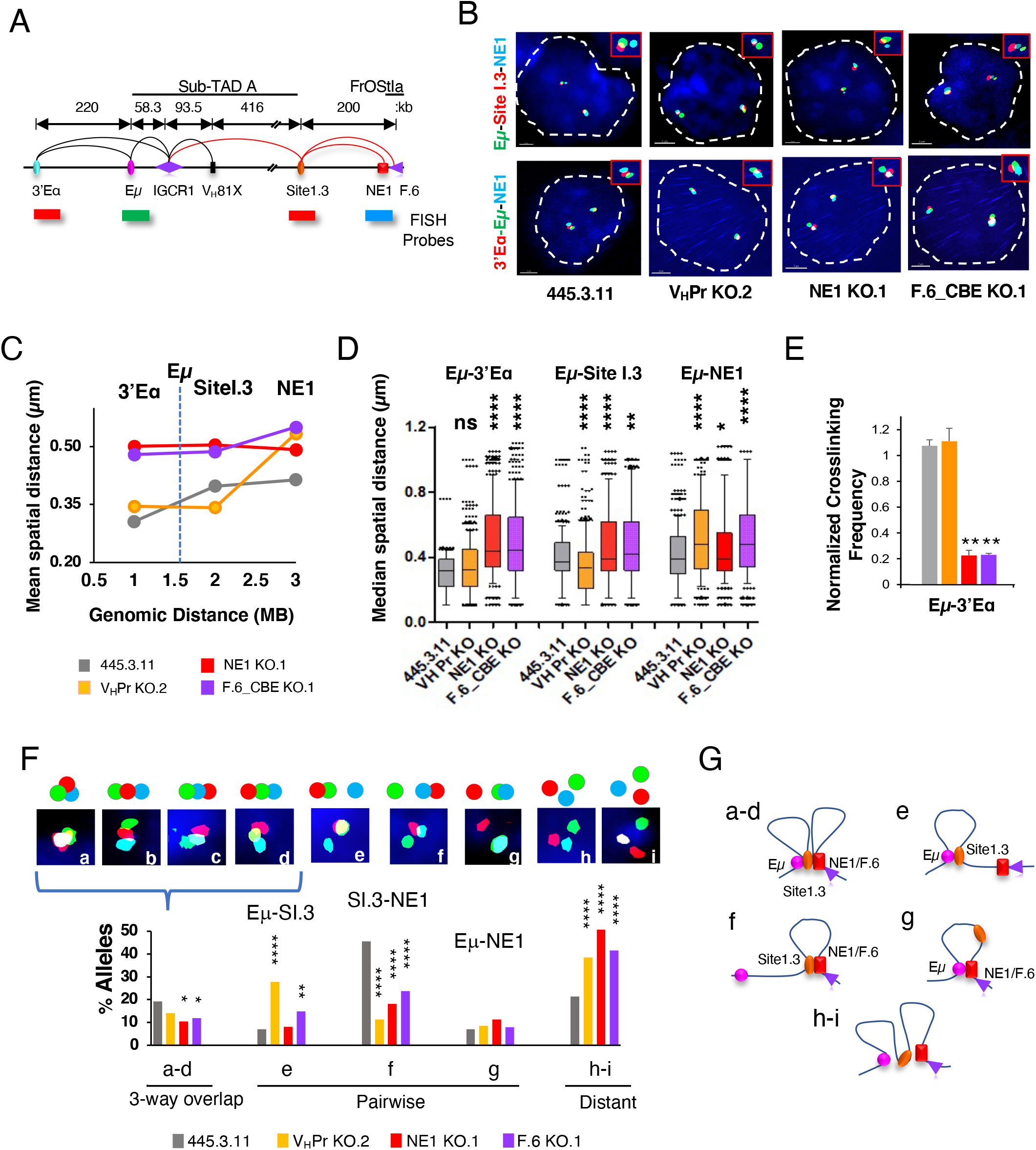
Transcriptional elements regulate locus compaction in Abl-t pro-B cells. **A)** Organization of the Igh locus from 3’Eα to FrOStIa and the position of FISH probes. Chromatin loops detected in 3C studies (red arcs; Figure 1D-F) and previous work (black arcs) (Kumar et al., 2013; Qiu *et al*., 2018; Zhang *et al*., 2019). Probes were labeled with AlexaFluor 488 (Eμ, green), AlexaFluor 555 (Site I.3 or 3’Eα, red), and AlexaFluor 647 (NE1, blue). **B)** Representative nuclei from 445.3.11 and KO lines were simultaneously hybridized with the indicated FISH probes (y-axis). **C)** Mean spatial distances were plotted as a function of genomic distance with lines indicating connectivity only. Vertical dashed line identifies the anchor probe position. Inter-probe distances for 404 alleles from two independent experiments for each probe combination were computed as described (Jhunjhunwala *et al*., 2008). **D)** Boxplots represent the distribution of spatial distances with the median shown. P values from Mann-Whitney U test. **E)** Normalized crosslinking frequencies for Eμ:3’Eα 3C looping interactions anchored at Eμ, analyzed in control and KO lines from at least from two independent experiments with SEMs. **F)** Quantitation of 3D probe configurations in three-color DNA FISH. (*Upper panel*) Nine probe configurations. (*Lower panel*) The frequency of each probe configuration. P values from Mann-Whitney U test. **G)** Representation of Igh alleles in spatial configurations (a-d, e, f, g, h-i).

To directly determine whether V_H_14-2 Pr-NE1/F.6 CBE assemble into a chromatin hub we analyzed FISH probe configurations using the Eμ (green), Site I.3 (V_H_14-2 Pr; red) and NE1 (blue) probes (fig. 3A). Molecular contacts between chromatin elements are indicated by superimposed FISH probes (≤ 0.3 μM). Segregation of probe contacts into structural categories allows identification of pure pairwise and three-way overlap interaction frequencies. Probes for Eμ, Site I.3 and NE1 were superimposed or overlapped as beads on a string in varying order in ∼20% of control alleles whereas the frequency of these probe configurations trended lower- or were significantly reduced in all three KO lines (fig. 3F, inset a-d). These findings demonstrate the formation of a chromatin hub that requires the presence of the V_H_14-2 Pr-, NE1- and F.6 CBE (fig. 3D, a-d). Significantly, the frequency of distantly spaced probes (≥ 0.31 μM) in which no overlap occurs greatly increased from ∼20% of alleles in controls to 40-50% in the KO lines demonstrating an overall destabilization of locus conformation by the loss of these elements (fig.3D inset h-i). The reduced presence of three-way interactions and the increased topological instability are consistent with reduced V->DJ recombination found for these KO lines. Finally, Eμ-Site I.3 looping increased upon V_H_14-2 KO indicating that this TE inhibits these interactions. We schematically represented an array of possible topological configurations including the Eμ-V_H_14-2-NE1/F.6 CBE chromatin hub that contribute to igh locus conformation (fig. 3G).

### V_H_ gene usage during V(D)J recombination is dependent on NE1 in mice

To assess the influence of NE1 on Igh repertoire formation and locus conformation in a physiological setting, we deleted 515 bp spanning NE1 using a genome editing strategy in mice (fig. 4A). NE1 deletion had little effect on B cell development as comparable proportions of Hardy fractions (B-C’, pro-B; D, pre-B; E, immature B; F, mature B) in the BM and marginal zone and follicular B cells in the spleen are detected in WT and NE1^-/-^ mice (Suppl. fig. 3A-E). To analyze the preselected Igh repertoire we FACS purified pro-B cells found in Hardy fractions B-C’ from WT and NE1-/-mice, and performed VDJ-seq, an unbiased assessment method for V_H_ segment usage in VDJ_H_ junctions (Suppl. fig. 3H) (Bolland *et al*., 2016; Chovanec et al., 2018). The overall profile for V_H_ gene usage in our WT samples compares well to previously reported findings (Bolland *et al*., 2016) (fig. 4B). In contrast, deletion of the NE1 element led to significantly reduced V_H_ gene usage in a localized domain referred to as the NE1 zone of influence (ZOI), extending from 5’ of V_H_81X (V_H_7183.2.3) to the V_H_J606 genes that mark the end of the intermediate genes and are located approximately halfway through subTAD B (fig 4C,D). V_H_ gene usage outside the ZOI was sporadically altered with increased usage near NE2 and NE3 implying widespread structure/function changes in NE1 deficient alleles (fig 4B,C).

**Figure 4.**
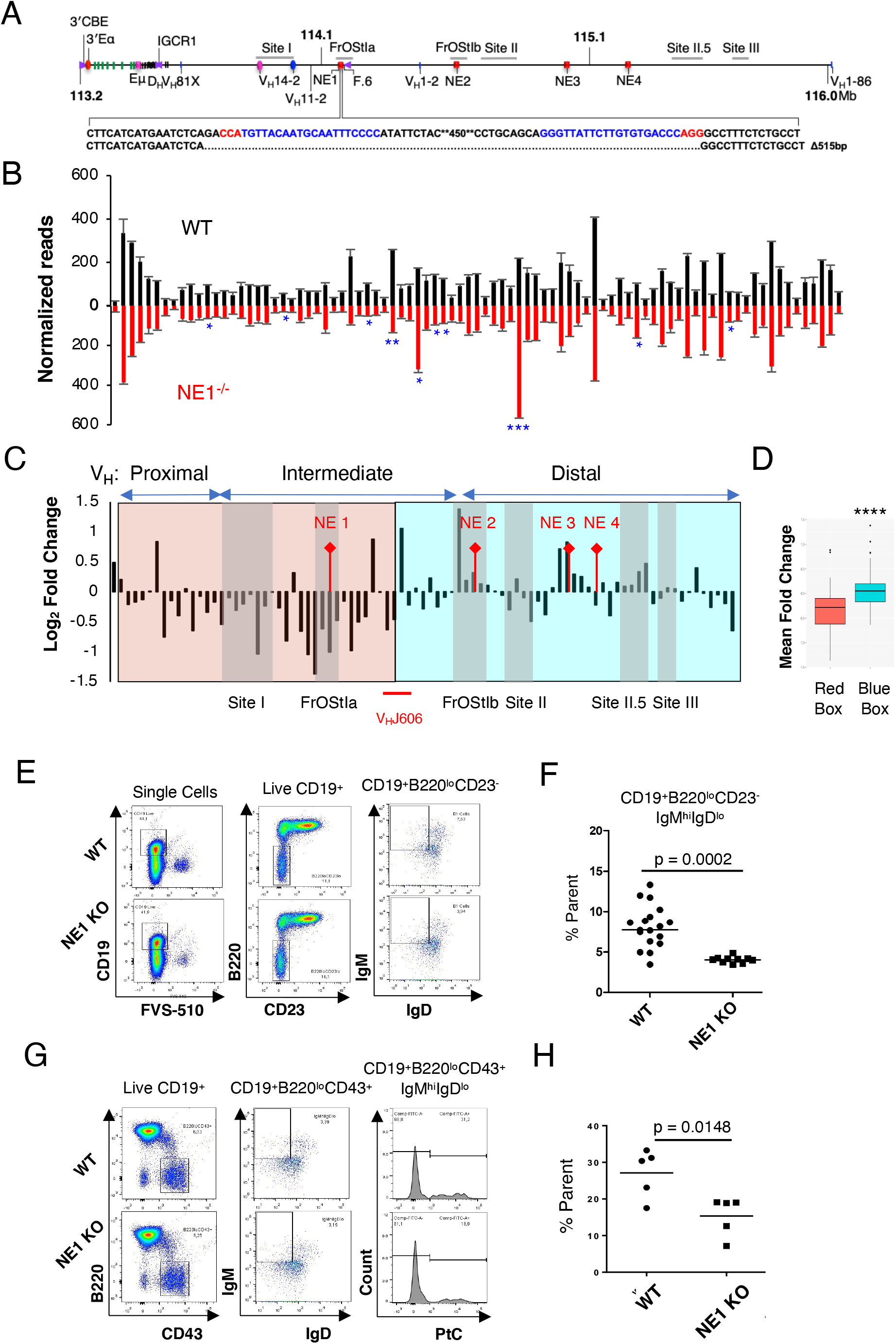
NE1 regulates regional V_H_ gene usage in mice. **A)** NE1 was deleted in mice using CRISPR/Cas9 genome editing. The location of the guide RNAs (blue) with PAM motifs (red) are indicated with the position and size of the NE1 deletion. **B**,**F**,**H)** P values from Student’s two-tailed t test. **B)** VDJ-seq analysis of gDNA from CD19^+^ pro-B cells from WT and NE1^-/-^ mice. Reads for V_H_ genes were normalized from two experiments, +/-SEM. **C)** Ratio of NE1^-/-^/WT normalized reads. Chromatin landmarks (grey rectangles) and V_H_J606 genes (red line) are indicated. **D)** VDJ-seq fold change (NE1^-/-^/WT) values were partitioned into the red and blue boxes **(C)** and analysed using the Mann Whitney Wilcoxon test. **E)** Representative flow cytometry plots of splenocytes gated for B1 B cells were assessed using FVS-510 and antibodies (CD19-PerCp, B220-APC Cy7, CD23-FITC, IgM-e450, IgD-AF700). **F**,**H)** The frequencies of B1 B cells were averaged. **F)** The frequency of B1 B cells (CD19^+^B220^lo^CD23^-^IgM^hi^IgD^lo^) was averaged. Each symbol represents results from one mouse. **G)** Representative flow cytometry plots of splenocytes gated for B1 B cells using FVS-510 and antibodies (CD19-PerCp, B220-APC Cy7, CD43-APC, IgM-e450, IgD AF700) and PtC liposome (FITC). **H)** The frequency of B1 B cells (CD19^+^B220^lo^CD43^+^IgM^hi^IgD^lo^PtC^+^) was averaged with each symbol representing the results from one mouse.

The NE1 ZOI encompasses the D_H_ proximal V_H_5 (V_H_7183) and V_H_2 (V_H_Q52) families and the small intermediate gene families including the V_H_6, V_H_11 and V_H_12 exons that are frequently expressed in B1 B cells (Prohaska et al., 2018; Yang et al., 2015). Normal proportions of peritoneal B1 B cells were detected in NE1^-/-^ as compared to WT mice (Suppl. fig. 3 F,G). Notably, splenic B1 B cells expressing surface CD19^+^B220^lo^CD23^-^IgM^hi^IgD^lo^ were significantly reduced in NE1^-/-^ relative to WT mice as are those specific for phosphatidylcholine (PtC) (Prohaska *et al*., 2018; Yang *et al*., 2015) are depleted in NE1-/-as compared to WT mice (fig. 4E-H). Thus, perturbation of the preselected Igh repertoire in pro-B cells has ramifications for the peripheral repertoire.

### NE1 is an Igh locus architectural element

To define the contribution of NE1 to locus architecture *in situ* Hi-C (Rao *et al*., 2014) data sets were generated from two biological replicates of Rag1^-/-^ and Rag1^-/-^NE1^-/-^ CD19^+^ pro-B cells that yielded a minimum of 1.3 billion read pairs and 0.72 billion contacts for each sample with a 10kb bin resolution (Suppl. Table 9). Rag deficient pro-B cells are used to ensure that the Igh locus has not undergone V(D)J recombination and are compared with those from mouse embryonic fibroblasts (MEF) cells in which the Igh locus is inactive (Di Giammartino et al., 2019). Pooled Hi-C samples recapitulated reported chromatin structures with high reproducibility scores and similar data quality (Suppl. Table 10) (Suppl. fig. 4A). The Igh 2.9 Mb TAD (yellow box), displays strong locus boundaries (green/blue arrows) and rarely interacts with other compartments (black brackets) as we previously observed for pro-B cells (Suppl. fig. 4A,B) (Montefiori *et al*., 2016). In contrast, the Igh locus in MEF is subsumed into an adjacent TAD and interacts with other compartments (Suppl. fig. 4A,B). Although the genomic flanking regions are quite structured, the Igh locus itself is highly self-interactive with few strong structural landmarks in pro-B cells (Suppl. fig. 4B). To identify pro-B cell specific interactions we created Hi-C difference heatmaps by subtracting MEF-from pro-B-derived interactions, a strategy used in our earlier studies (Montefiori *et al*., 2016). The Rag1^-/-^ minus MEF difference map reveals stripes (blue arrowheads) originating from the 3’RR and IGCR1 as previously noted (Benner et al., 2015; Vian et al., 2018) that are absent in MEF (Suppl. fig. 4A,B). Nested self-interacting domains (vertical dashed lines) spanning hundreds of kb along the locus are detected in the pro-B cell difference map and have been schematically summarized (fig. 5Bi, iii). We also observe a novel series of stripes (black arrowheads) some of which are clustered, and flames extending in a 5’->3’ direction from the corners of multiple self-interacting domains (fig. 5Bi). Several of these stripes meet off the diagonal to form “dots” indicating the presence of IGCR1-NE1 and IGCR1-NE2 (marked a and b), NE1-NE2 (marked c), and V_H_J606 genes-NE2 (marked d) anchored loops indicating that loop domains anchored by NE-NE and NE-Pr contacts contribute to Igh locus conformation (fig. 5B iii). Architectural stripes, are thought to form via cohesin mediated loop extrusion (Fudenberg *et al*., 2016; Vian *et al*., 2018) and to facilitate V(D)J joining in pro-B cells (Hu et al., 2015).

**Figure 5.**
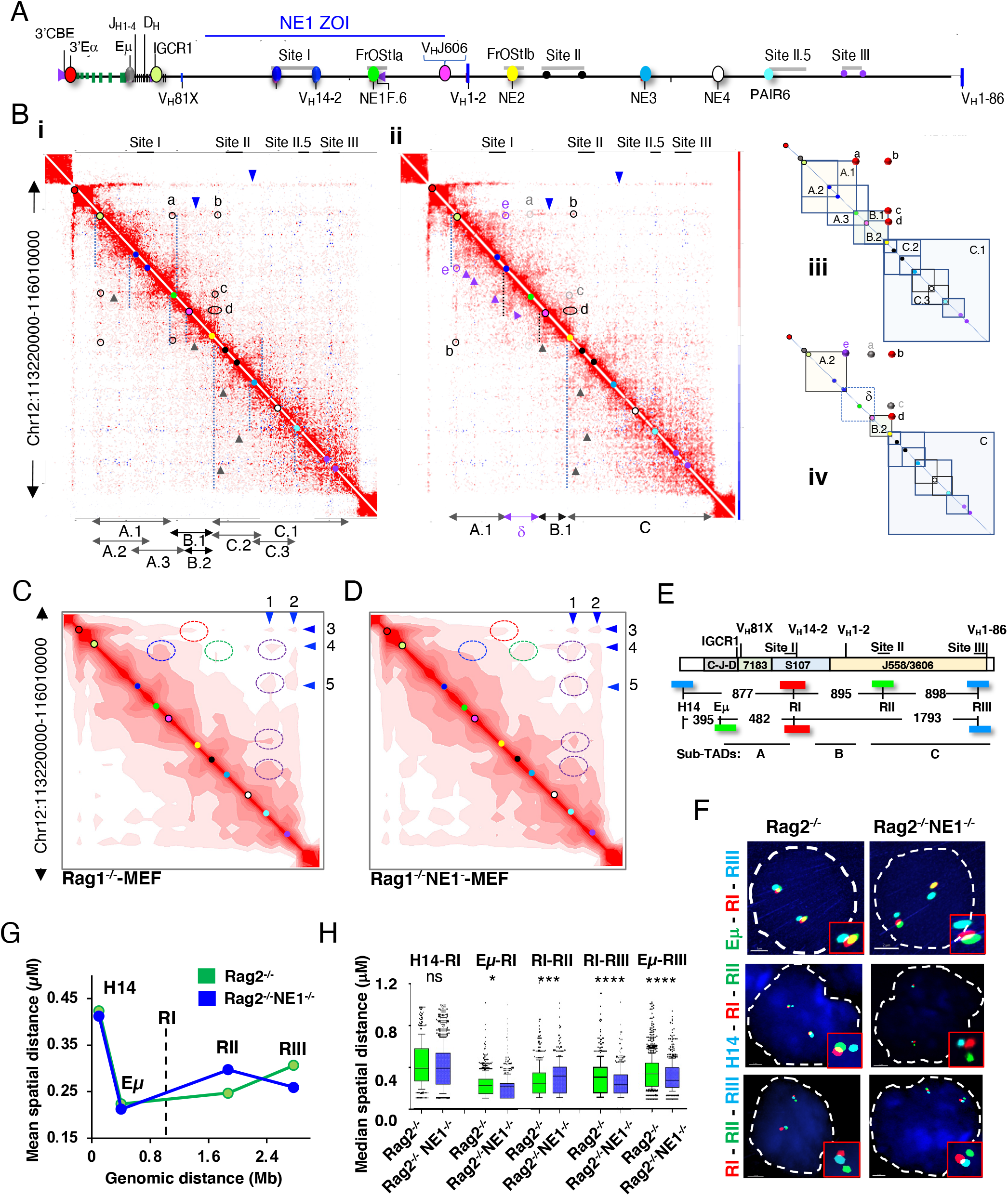
NE1 modulates Igh locus topology and compaction in pro-B cells. **A)** Schematic of the Igh locus. **B-D)** Regulatory elements identified in **(A)** have been arrayed along the Hi-C diagonal as color coded reference dots. Hi-C difference heatmaps at 10kb resolution from purified CD19^+^ pro-B cells from Rag1^-/-^ **(i)** and Rag1^-/-^NE1^-/-^ **(ii)** mice. Data were KR normalized and then quantile normalized using a random model. Nested loop domains (dashed vertical lines), stripes (3’->5’, blue triangles, 5’->3’ black triangles), dots (black circles), lost dots (gray circles). Rag1^-/-^NE1^-/-^ specific stripes (purple triangles) and dots (purple circles). **iii, iv)** Schematic of nested loop domains found in Rag1^-/-^ (**iii)**, and Rag1^-/-^NE1^-/-^ **(iv)** from Hi-C difference heatmaps. **C**,**D)** Hi-C difference heatmap maps at 100kb resolution. Stripes (blue triangles). **E)** Genomic organization of the Igh locus with BAC probes used in three independent FISH experiments. **F)** Representative nuclei from Rag2^-/-^ and Rag2^-/-^NE1^-/-^ pro-B cells with labeled FISH probes hybridized to fixed cells. **G)** Average inter-probe distances displayed as a function of genomic distance and lines indicate connectivity only. Vertical dashed line indicates the anchor position. **H)** Boxplots show the distribution of spatial distances with medians between probes and anchor R1. P values from Mann-Whitney U test.

The structural profile of the Igh locus was substantially altered in Rag1^-/-^NE1^-/-^ derived Hi-C difference maps (fig. 5B i,ii). Many loops involving NE1 were abolished including those between IGCR1-NE1 (marked a, grey) and NE1-NE2 (marked c, grey), whereas those involving the NE2 anchor such as IGCR1-NE2 (marked b), and V_H_J606 genes-NE2 (marked d) are retained in the Rag1^-/-^NE1^-/-^ pro-B cells (fig. 5B ii, iv). Deletion of NE1 also led to the new appearance of IGCR1-Site I (marked e, purple circle) interactions, the formation of the δ loop (dashed box) configured by the association of Site I-V_H_J606 genes and a series of new stripes and chevrons (purple arrowheads) (fig. 5B ii, iv). Impairment of Igh locus structure conferred by NE1 deletion extends from IGCR1 to the J606 genes and maps well to the NE1 ZOI, thus, linking Igh spatial architecture with V_H_ gene utilization.

### NE1 constrains loop extrusion

To assess the higher-order conformation of the Igh locus we displayed the Hi-C data at 100kb resolution and note several important differences between the Rag1^-/-^ and Rag1^-/-^NE1^-/-^ difference heatmaps (fig. 5C,D). First, IGCR1:NE1 contacts (blue dashed circles) that are evident in Rag1^-/-^ are absent in Rag1^-/-^NE1^-/-^ difference maps in accord with findings in the high resolution heatmaps (fig. 5C,D). Second, NE2 sits at the pivot point between sub-TADs A-B and C in maps for both genotypes suggesting an important architectural function for this enhancer (fig. 5C,D). Finally, we note at least four long stripes (marked by blue triangles) in Hi-C maps in both Rag1^-/-^ and Rag1^-/-^NE1^-/-^ Hi-C maps. Stripes that originate at PAIR6 (1) and Site III (2) extend in the 5’ to 3’ direction whereas those that initiate at 3’Eα (3) and IGCR1 (4) extend in the 3’ to 5’ direction (fig. 5C,D). Stripes initiating at PAIR6 (1), Site III (2) and IGCR1 (4), appear punctuated with discrete stopping points that could give rise to loop domains. For example, PAIR6 interactions with NE3, NE2, Site I, and IGCR1 (purple ovals) are punctate and more frequent in the Rag1^-/-^NE1^-/-^ difference maps (fig. 5C,D). Hence, NE1 may function to constrain loop extrusion and thereby reduce the long range interactions between 5’ and 3’ ends of the locus that might confer locus compaction.

### NE1 deletion leads to Igh locus hyper-contraction

To determine the contribution of NE1 to the higher order conformation of the Igh locus we compared the average- and the median-spatial distances between FISH probes (Giorgetti and Heard, 2016) in Rag2^-/-^ and Rag2^-/-^NE1^-/-^ pro-B cells. The bacterial artificial chromosome (BAC) probes H14, located outside the Igh locus, Eμ-IGCR1 (referred to as Eμ) probe spanning the RC and D_H_J_H_ elements and RI, RII and RIII probes, corresponding to Sites I, II and III, were hybridized in pro-B cells (fig. 1G, 5E,F) (Montefiori *et al*., 2016). We used probe RI, that overlaps the intermediate V_H_ gene segments, as an anchor to determine the spatial distances with all other probes. Although spread over a large genomic distance the average spatial distance separating probes RI from Eμ, RII and RIII were similar whereas spatial distances to the H14 probe outside the locus were large indicating that most V_H_ genes are at similar distances to the RC in Rag2^-/-^ pro-B cells, as previously observed (fig 5G) (Jhunjhunwala *et al*., 2008). In contrast, closer spatial distances of between the anchor probe RI to RIII in Rag2^-/-^NE1^-/-^ pro-B cells reflecting a higher degree of compaction between Eμ and the distal V_H_ gene segments relative to the control (fig 5G). This conclusion is also supported by the decreased spatial distances for probes Eμ to RIII (fig. 5H). Thus, increased architectural stripe length upon NE1 deletion is correlated with increased locus compaction.

### NE1 coordinates an enhancer network

The Hi-C analyses indicated that NE1 interacts with NE2. To explore the scope NE interactions and their impact on Igh locus topology we reduced Hi-C dimensionality by deriving virtual 4C datasets. Using NE1, Eμ and IGCR1 viewpoints we identified high frequency interactions defined as in the top 15 (star) or 25 (circle) percent of all contacts in both biological replicates for Rag1^-/-^ (black symbols) and Rag1^-/-^NE1^-/-^ (red symbols) pro-B cells (fig. 6A). NE1 was highly interactive with its own flanking region, Eμ, NE2, NE3, and NE4 in Rag1^-/-^ pro-B cells (fig. 6A). Although NE3 associates with NE1 its interaction frequency did not pass the threshold for high frequency contacts in both replicates (fig. 6A). Deletion of NE1 (red dashed line) led to loss of contact with all other NEs and with Eμ and the emergence of two new peaks (red arrows) of contact at Site I.3 and NE2 in Rag1^-/-^NE1^-/-^ pro-B cells and suggests that NE1 coordinates an enhancer network (fig. 6A).

**Figure 6:**
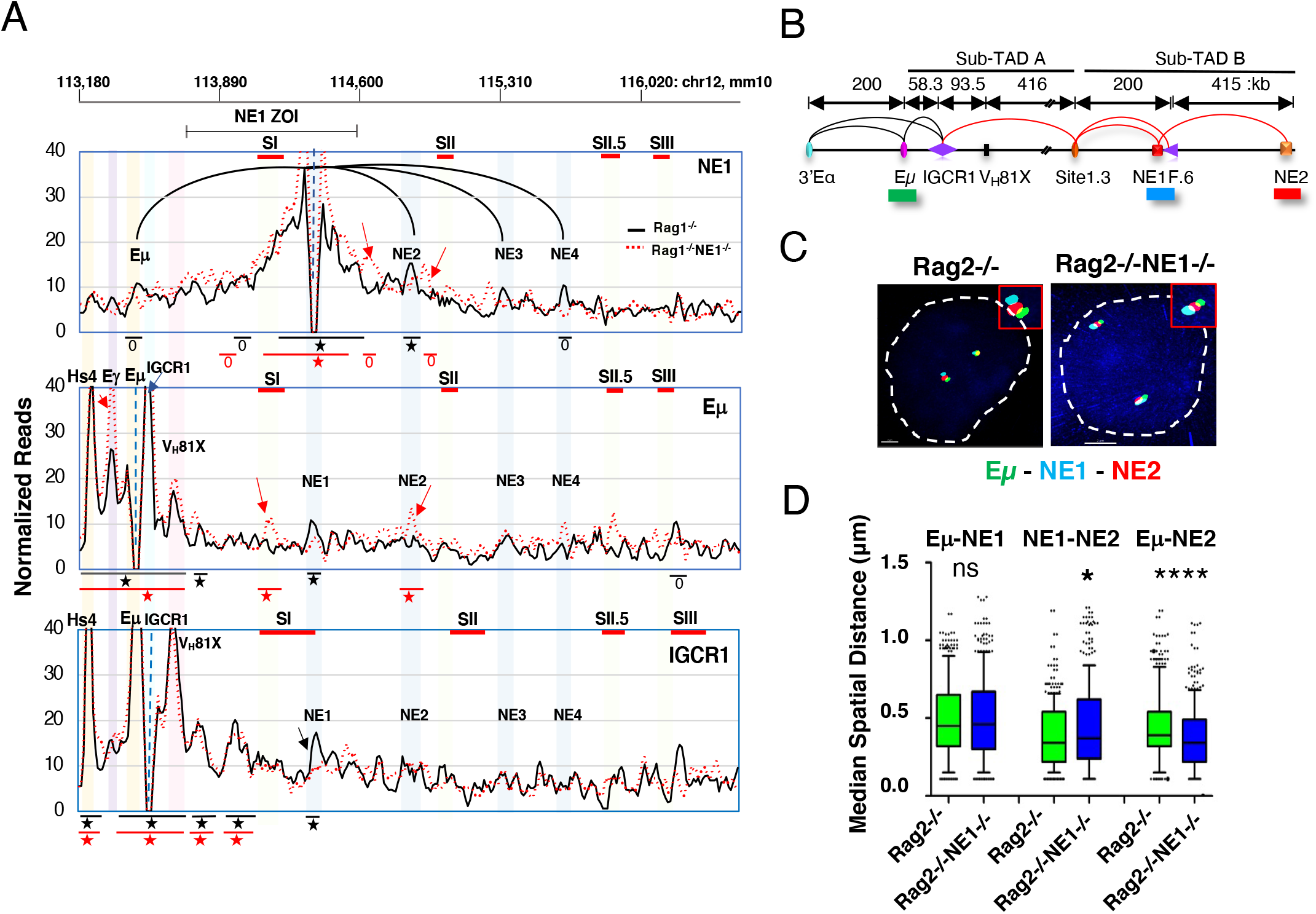
A NE interactome identified. **A)** Virtual 4C interactions were extracted from KR normalized Hi-C data sets from Rag1^-/-^ (black lines) and Rag1^-/-^NE1^-/-^ (dashed red lines) CD19^+^ pro-B cells. *Top*: Genomic coordinates (mm10) with the NE1 ZOI and Sites (S) I, II, II.5 and III. 4C viewpoints (dashed vertical line) in a 30 kb running window analysis with 10 kb steps from merged biological replicates with NE1 interactions (black arcs) shown. Regions marked by stars (top 15%) or circles (top 25%) of locus wide interactions. New peaks in Rag1^-/-^NE1^-/-^ pro-B cells (red arrows) and lost peaks (black arrows) identified. **B)** Diagram of the Igh locus with genomic distances and FISH probe indicated. **C)** Representative nuclei from Rag2^-/-^ and Rag2^-/-^NE1^-/-^ CD19^+^ pro-B cells. Short FISH probes, Alexa Fluor 488 (Eμ, green), and Alexa Fluor 647 (NE1, blue), Alexa Fluor 555 (NE2, red) were hybridized simultaneously to fixed cells in duplicate. **D)** Boxplots showing the median and distribution of spatial distances between two probes. P values from Mann-Whitney U test.

Next, we examined the Eμ viewpoint and confirmed interaction with 3’Eα hs4, Eγ, and IGCR1 as previously reported (Dai *et al*., 2021; Jain *et al*., 2018; Predeus et al., 2014; Qiu *et al*., 2018) and with NE1 (fig. 6A). Eμ:NE1 contacts were lost and new or increased Eμ interactions (red arrows) with Eγ, Site I.3, and NE2 were detected in Rag1^-/-^NE1^-/-^ pro-B cells suggesting that NE1 competes with Eγ and NE2 for interaction with Eμ (fig. 6A). Finally, analysis of the IGCR1 viewpoint revealed interactions with 3’Eα hs4, Eμ, V_H_81X and NE1 but not with Eγ or with other NEs indicating that IGCR1 could bridge Eμ to NE1 (fig. 6A). We infer that NE1 may integrate contacts with Eμ, IGCR1 and the distal NEs.

To independently confirm that Eμ interacts with NE2 upon NE1 deletion we performed FISH and examined pairwise inter-probe spatial distance distributions using short FISH probes for Eμ (green), NE1 (blue), and NE2 (red) in Rag2^-/-^ and Rag2^-/-^NE1^-/-^ pro-B cells (fig. 6 B,C). Indeed the median spatial distances for Eμ:NE2 were significantly reduced and NE1:NE2 probe distances were markedly increased in Rag2^-/-^NE1^-/-^ pro-B cells, consistent with the virtual 4C profile (fig. 6D). Hence, NE2 becomes the preferred Eμ interaction partner in the absence of NE1 indicating that Igh enhancers compete for interaction partners.

### NE1 is a negative regulator of V_H_ gene expression

To assess the enhancer function of NE1 we evaluated transcriptional activity for a set of sense and anti-sense intergenic Igh index genes (Bolland et al., 2004; Oudinet et al., 2020) and found varying levels of NE1 dependency. As in the 445.11 NE1 KO lines, the V_H_14-2 GLT was significantly reduced in Rag2^-/-^NE1^-/-^ pro-B cells indicating NE1 dependency (Suppl fig. 7A,B). To more broadly assess V_H_ GLT levels we performed RNA-seq and found significantly elevated expression of annotated V_H_ genes in Rag2^-/-^NE1^-/-^ pro-B cells as compared to controls (fig. 7D-F). In contrast, gene expression was similar for both genotypes in a 1 Mb region containing 53 expressed genes downstream of the Igh locus (fig. 7F). One interpretation of these findings is that NE1 is a negative modulator of V_H_ GLT expression. Another possibility is that alteration of Igh locus conformation associated with NE1 deletion led to a more permissive transcriptional environment. Indeed, the increased association of distal Igh regions with Eμ, detected in our FISH studies, may provide an enhanced transcriptional environment (fig. 5H, 6E). Nevertheless, how can we reconcile the increase of V_H_ gene expression and decreased V_H_ gene usage in the NE1 ZOI?

**Figure 7.**
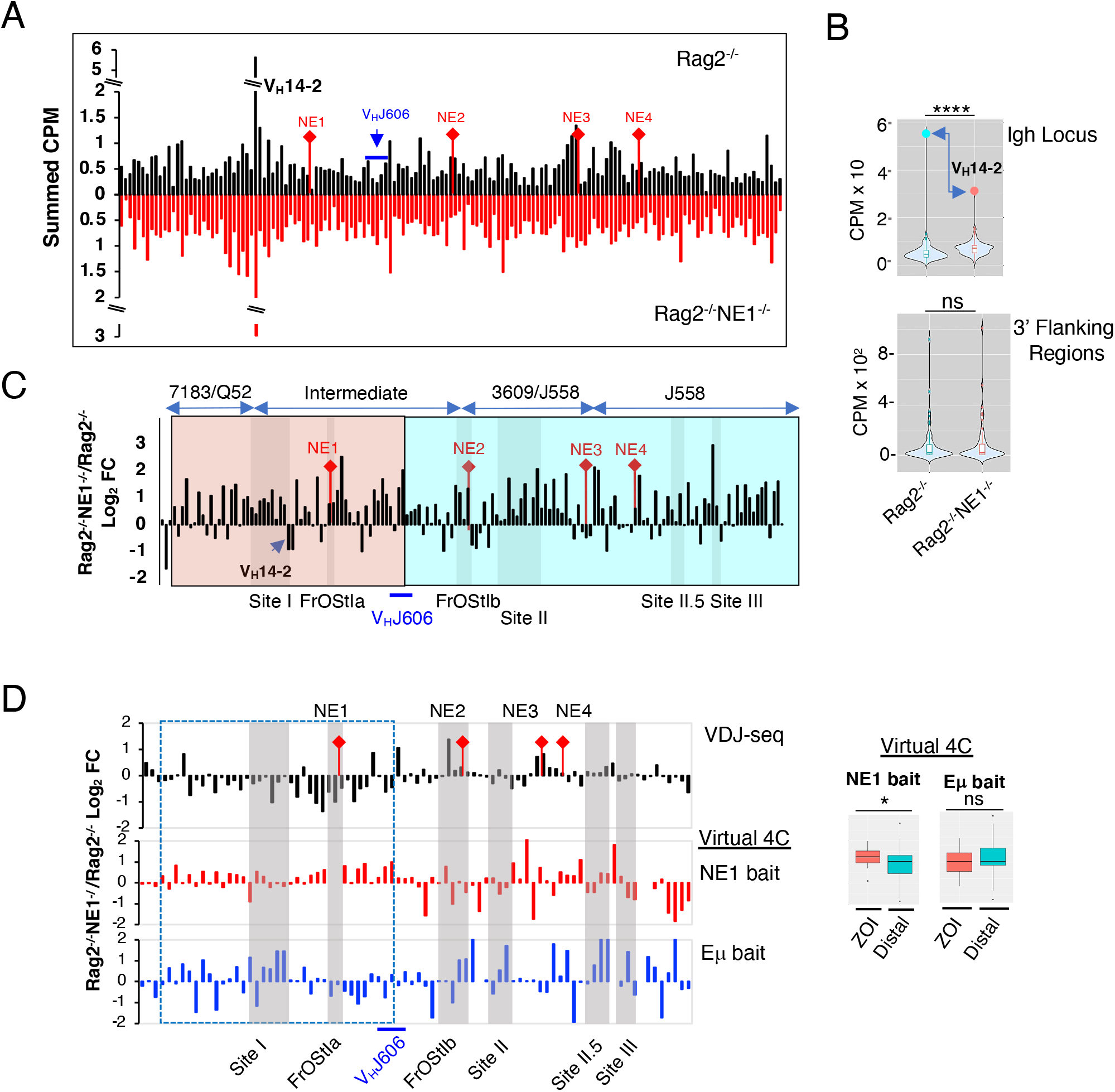
NE1 regulates V_H_ transcription locus-wide. **A)** Normalized RNA-seq data for annotated V_H_ genes from Rag2^-/-^ and Rag2^-/-^NE1^-/-^ pro-B cells. Three independent samples were summed for each genotype. Individual V_H_ genes were included when there was >0.3 cpm in at least one genotype. **B)** Violin plots compare the distribution of cpm values from two genotypes. P value from Mann Whitney Wilcoxon test. **C)** Ratio of Rag2^-/-^NE1^-/-^/Rag2^-/-^ RNA-seq cpm expressed as log2 fold change (FC). NE1 ZOI (pink box) and Igh distal region (blue box) defined for V_H_ gene usage in Figure 4C. **D)** Log2 fold change (FC) of Rag1^-/-^NE1^-/-^/Rag1^-/-^ normalized VDJ-seq and the virtual 4C reads for the NE1 and Eμ viewpoints in pro-B cells. Virtual 4C contacts were aligned to the same 10 kb bins that contain V_H_ genes analyzed in the VDJ-seq study. NE1 ZOI indicated by the dashed blue box. P values from Mann Whitney Wilcoxon test.

### NE1 dependent V_H_ gene usage is correlated with chromatin conformation

To determine whether the NE1 ZOI chromatin landscape was uniquely affected by NE1 deletion. we compared the V_H_ gene usage profile (top panel) to that of normalized virtual 4C contacts for the NE1- and Eμ-viewpoints (fig. 7G). NE1 regional contacts were significantly elevated in the ZOI and randomly altered outside of this region whereas there was no discernable pattern for Eμ chromatin contacts in Rag1^-/-^NE1^-/-^ pro-B cells (fig. 7G, right panel). We propose that NE1 may underpin an architectural structure that promotes ZOI V_H_ gene usage by excluding extraneous chromatin contacts. NE1 deletion also leads to increased locus contraction as noted by the increased proximity of distal V_H_ genes to Eμ (fig. 5H, 6E). Therefore, it is possible that NE1 modulates V_H_ gene expression locus-wide by constraining locus compaction. Our studies imply that chromatin conformational states select for transcriptional versus recombinational potential.

## DISCUSSION

We report a major role for Igh associated NEs and TEs in configuring locus topology and in V(D)J recombination. Our findings identify NE1, the V_H_14-2 Pr and F.6 CBE as regulators of V->DJ recombination potential and locus conformation and reveal a structure/function relationship in Abl-t pro-B cell lines. To broaden this analysis we deleted NE1 in mice and found that NE1 loss led to regionally reduced V_H_ gene usage and to a dramatic reduction of splenic B1 B cells thus, linking the quality of the pre-selected repertoire to the shape of the mature Ig repertoire in the periphery.

Chromatin conformational analysis permitted mapping of pro-B cell specific interactions and highlighted NE1 and NE2 as pivotal structural elements in Igh TAD organization. Our data provide strong evidence that architectural “stripes” initiate from the 3’ and 5’ ends of the locus in pro-B cells. Stripes form when one subunit of cohesin stalls near a strong CTCF loop anchor and the second subunit progressively extrudes the chromatin loop to form a multiplicity of contacts and that ultimately brings the distal end of the locus into spatial proximity to the stall site (Vian *et al*., 2018). Extrusion initiated at the 3’ end of the Igh locus could bring intermediate V_H_ genes into physical proximity with the RC and has been used to describe the trajectory of RAG scanning model (Dai *et al*., 2021). Although stripes originating at the 5’ end of the locus would have the wrong polarity to support RAG scanning this extrusion trajectory could establish E-E-Pr anchored loops and loop domains that link VH genes in subTAD C to IGCR1 and the RC.

Convergent CBE and other elements including enhancers can substantially block extrusion or could permit a series of soft stops and read-throughs that would form loops of different lengths and different final endpoints (Dekker and Mirny, 2016; Rowley and Corces, 2018). Indeed, Igh stripes show a series of punctate stops that align with loop anchor sites including NE1, NE2, NE3 and Site I. NE1 appears to block both upstream and downstream extrusion and thereby limit the length of these stripes. NE1 deletion relieves the block and allows robust extrusion to extend most prominently from the 5’ end of the locus and facilitates interactions with the downstream RC. Igh locus contraction juxtaposes distal V_H_ genes to proximal D_H_ segments to promote V(D)J recombination (Fuxa *et al*., 2004; Jhunjhunwala *et al*., 2008; Kosak *et al*., 2002; Roldan *et al*., 2005; Sayegh *et al*., 2005). Although loop extrusion has been proposed to generate locus compaction (Hill *et al*., 2020) direct evidence has been lacking. In addition to causing extended extrusion along the entire length of the Igh locus we found increased locus compaction in NE1 deficient pro-B cells, revealing a direct link between these processes.

NE1 has enhancer activity as the transcriptional potential of several V_H_ genes and AS intergenic transcripts was impaired in Abl-t NE1 KO pro-B cell lines. Contrary to expectations, expression of annotated V_H_ GLTs increased locus-wide in NE1 deficient primary pro-B cells indicating that this NE is a negative regulator of V_H_ expression and that another potent enhancer(s) gains access to V_H_ Prs locus-wide. Many enhancers regulate transcription with their cognate Prs via spatial proximity created by long range interactions (Furlong and Levine, 2018) or by co-compartmentalization into transcriptional condensates (Cho et al., 2018; Shrinivas et al., 2019). Loss of NE1 produced significant regional and locus-wide topological perturbations that served to increase overall locus contraction, increase proximity of V_H_ Prs to Eμ and create a pretext for elevated V_H_ transcription.

NE1 affected transcription locus-wide yet it controls rearrangement over a much smaller region in effect segregating these functions. Likewise, recent studies identified E88 is a structural organizer of the Ig*k* locus locus (Barajas-Mora et al., 2019). However, E88 is distinct from NE1 as it impacted transcription over a relatively small area and yet regulates V_H_ gene rearrangement over a much broader area (Barajas-Mora *et al*., 2019). This difference may be related to the structural roles NE1 plays as 1) a stripe origin running in the 5’ to 3’ direction and as 2) a block to extrusion that in turn modulates locus compaction. Our studies have distinguished chromatin conformational states that select for broad V_H_ gene transcriptional competence on the one hand or V(D)J recombinational potential in the NE1 ZOI on the other.

## ACKNOWLEDGEMENTS

This work was supported by grants from the NIH to A.L.K. and A.J.F. (RO1AI121286) and to

A.L.K. (R21AI133050). The authors declare that they have no competing financial interests.

## AUTHOR CONTRIBUTIONS

Conceptualization, A.L.K., A.J.F., Methodology, X.Q., R.W., H.F., J.L., E.K., A.J.F., A.L.K., Validation, R.W., J.F.C., Investigation, K.H.B., X.Q., S.N., S.P., R.W., J.F.C., E.K., J.X., Formal Analysis, H.F., X.L., J.L., S.N., Resources, A.L.K., A.J.F., N.B., J.L., Writing-Original Draft, A.L.K., Writing-Review and Editing, A.L.K., A.J.F., J.L., N.B., S.P., K.H.B., Visualization, H.F., X.L., S.N., K.H.B., J.F.C., J.L., A.L.K., Supervision, Funding Acquisition, A.L.K. and A.J.F., Project Administration, A.L.K.

