## Supplementary material for "An Igh novel enhancer modulates antigen receptor diversity by determining locus conformation": Suppl. Figures, Methods and Tables

#### Supplementary Information

#### METHODS

##### Mice, cell lines and cell culture

C57BL/6 (WT) or Rag deficient mice on the C57BL/6 background were purchased from Jackson Laboratories or maintained in colonies at the University of Illinois College of Medicine, or Scripps Research. All procedures involving mice were approved by the Institutional Animal Care Committee of the University of Illinois College of Medicine, and the Scripps Research Institute, in accordance with protocols approved by the UIC and Scripps Research Institute Institutional Animal Care and Use Committees. Rag deficient CD19+ pro-B cells were isolated from BM using anti-CD19 coupled magnetic beads (Miltenyi) and cultured in the presence of IL7 (1% vol/vol supernatant of a J558L cell line stably expressing IL7) for 4 days. The Abelson-MuLV transformed (Abl-t) pro-B cell line, 445.3 (Rag1-/-) on the C57Bl/6 background was kindly provided by Dr. B. Sleckman (University of Alabama at Birmingham) (Kumar *et al.*, 2013). A newly derived subclone, 445.3.11 from the Abl-t 445.3 line were cultured in RPMI 1640 (Cellgro), 10% (v/v) FBS, 4mM glutamine (Gibco), 1mM sodium pyruvate (Gibco), 1X nonessential amino acid (Gibco), 5000 units/ml Penicillin and 5000 mg/ml Streptomycin (Gibco), 50 mM $\beta$-mercaptoethanol (Sigma) and maintained at approximately 5x10e5 cells/ml. Splenic T cells were enriched using Mouse T Cell Enrichment Columns (MTCC-5; R&D Systems) and cultured at a density of $5X10^5$ to $1X10^6$, stimulated in RPMI 1640 and glutamine (4 mM) and penicillin- streptomycin supplemented with FCS (10% v/v), and activated with Con A (5 ng/ml; 15324505; MP Biomedicals).

##### Flow cytometry

Flow cytometry analyses were performed using live pro-B cells ($5x10^{-5}$) washed in PBS plus 2% FCS, and stained with antibodies (CD19-PerCp, B220-APC Cy7, CD93 PE Cy7, CD43-APC, CD2-PE, IgM-e450) by gating for Fixable Viability Stain 510 (FVS510) (BD Biosciences) and/or forward- and side scatter on a CyAn ADP with Summit software (Becton Coulter, Indianapolis, IN), or an Attune (Invitrogen, CA) with Flowjo software (Flowjo LLC, OR). Assessment of splenic marginal zone (MZ) and follicular (Fo) B cells from mice was performed by flow cytometry. Splenocytes were isolated and the live B cells were enumerated by using the viability stain FVS-510, and the antibodies B220-APC Cy7, CD21-FITC, CD23-biotin and streptavidin-APC. Splenocytes were isolated and the live B1 B cell component analyzed by treating with viability stain FVS-510, and with antibodies to CD19-PerCp, B220-APC Cy7, CD23-FITC, anti-IgM-e450, IgD AF700, CD5-PE or with CD19-PerCp, B220-APC Cy7, CD43-APC, IgM-e450, IgD AF700 and PtC liposomes (FITC) (a gift from N. Baumgarth, UC Davis) and analyzed on a BD LSRFortessa with gating as described (Baumgarth, 2004; 2011) with FlowJo software. All antibodies and reagents are listed in (Suppl. Table 1).

**Quantitative RT-PCR and ChIP**

RNA extraction was performed from 2-3x$10^6$ cells using TRIzol (Life Technologies) according to manufacturer instructions. RNA samples were further treated with the DNase I kit (Invitrogen) to remove contaminating DNA per the manufacturer's instruction. First strand cDNA synthesis was done with RNA (1-4 μg) and the SuperScript III Reverse Transcriptase kit (Invitrogen) according to the manufacturer's instructions. Quantitative (q) real-time RT-PCR was performed using primers (Suppl. Table 2) and Fast SYBR Green PCR Mix (Applied Biosystems) and Real-Time PCR System viiA7 (Applied Biosystems) as described (Wuerffel et al., 2007) except that primers for 18S rRNA were used (Rhinn et al., 2008) to normalize samples. Semi-quantitative RT-PCR assays for $V_H$J558 transcripts were carried out using Platinum Taq DNA polymerase (Invitrogen) (25 μl) at 94°C, 30 s for 1x; 94°C 30 s, 60°C 30 s, 72°C 30 s, for 29x, 31x and 33x and 7 μl were analyzed by gel electrophoresis. The 18S loading control was assessed by qRT-PCR. Semi-quantitative RT-PCR assays were carried out for the $V_H$81X and $V_H$2-5 genes using the same conditions as for VHJ558 except that the PCR products were harvested at 32 cycles and for 18S rRNA at 28 cycles. All results represent the average of at least three independent experiments from 3-5 mice for each genotype or 2 biallelic identical 445.3.11 KO subclones. Each sample was assayed in triplicate and SEMs were calculated. ChIP assays for CTCF binding were performed with anti-CTCF antisera as described previously (Suppl. Table 1) (Feldman et al., 2017).

**CRISPR-Cas9 mediated genomic editing of cell lines and mice**

The optimal gRNA sequence closest to the genomic targets were identified using the http://crispr.mit.edu design tool and gRNAs were cloned into the pX330 vector at the BbsI site (Suppl. Table 3) (Ran et al., 2013). GRNA efficiency was tested in HEK293T cells using previously described protocol (Mashiko et al., 2013). The Abl-t 445.3.11 cell line was co-transfected with gRNAs plasmid constructs (1 μg/each) together with a pmax GFP plasmid (1 μg) (Lonza) using the Amaxa Cell Line Nucleofector Kit (Lonza) (Y001 program) and a Nucleofector II (model AAD-1001N) according to the manufacturer's instructions. Cells were allowed to recover for 48 h and then GFP+ cells were purified by FACS (MoFlo Astrios) supported by Summit software (Beckman Coulter, Indianapolis, IN). Cells were submitted to limiting dilution, subclones were expanded for 10–14 days and gDNA was harvested using the alkaline lysis method. Subclones were screened for insertion/deletions (indels) by PCR using Platinum Taq DNA polymerase (Invitrogen) and target specific primers at 94°C, 30 s for 1 cycle; 94°C 30 s, 58-66°C 30 s, 72°C 30-60 s for 35 cycles; 72°C 5 min for 1 cycle (Suppl. Table 4). PCR products were examined by Sanger sequencing to authenticate CRISPR/Cas9 indels.

To construct NE1-/- mice, Cas9 mRNA and sgRNAs NE1 g1 and NE1 g2 were injected into pronuclei of mouse zygotes at the Scripps Research Institute Mouse Genetics Core facility (Suppl. Table 3) (Mashiko *et al.*, 2013). The $NE1^{-/-}$ mouse contains a 515 bp deletion (chr12:114182511-114183025, mm10) spanning NE1.

**V(D)J recombination assays in Abl-t cells**

Abl-t 445.3.11 Rag1-/- cells were transfected with pMSCV-IRES-Bsr-Rag1 plasmid (a gift from Dr. D. Schatz, Yale University) using an Amaxa Cell Line Nucleofector Kit (Lonza) (Y001 program) and a Nucleofector II (Lonza, model AAD-1001N) according to the manufacturer's instructions. Stable transfectants were selected in 20 μg/ml blasticidin for 10-14 days and subsequently maintained in 10 μg/ml blasticidin. Cells ($1X10^7$) were treated with STI-571 (2.5 μM) for 48 hours and gDNA and RNA were isolated. RNA was prepared from cells ($9x10^6$)

using TRIzol (Life Technologies) following the manufacturer's instructions and gDNA was prepared from cells (1x$10^6$) by spooling. $D_H$->$J_H$ rearrangement was assayed using gDNA (50 ng) with DFL16.1F and JH1R primers and the Fast SYBR green master mix (Applied Biosystems) with qPCR of the Mb1 gene as the loading control. V->DJ recombination was assayed using cDNA (2 µl) in Fast SYBR Green PCR Mix (10 µl reaction volume) in a viiA7 system (Applied Biosystems) with 18S RNA as a loading control. Primers are listed in Suppl. Tables 2, 5.

**DNA fluorescent in situ hybridization (FISH)**

DNA probes for fluorescent in situ hybridization (FISH) were prepared from locus specific BACs or short probes. All genomic coordinates are chr12 (mm10). The BACs probes were RI (113813346-113989276) (BAC 373N4), RII (114705813-114886430) (BAC 70F21), RIII (115539177-115773011) (BAC 368C22 or 230L2) (Montefiori *et al.*, 2016)and H14 (112907511 – 113142759) located ~44 kb 3′ of HS7 of the 3′RR (BAC RP23-201H14) (Gerasimova et al., 2015). Short probes (4.2-7.5kb) were Eµ (113425893–113430719), Site I.3 (113987323–113994353), NE1 (114184075–114191532) and NE2 (114600831–114606206) were generated by PCR using gDNA with primers listed in (Suppl. Table 6). Probes were labeled by nick translation in the presence of Alexa Fluor 555 (red) and 488 (green) and Alexa Fluor 647 (blue). Labeled probes were hybridized with fixed cells and FISH was performed using a Nikon W1 dual CAM spinning disk confocal microscope (Center for Advanced Microscopy and Nikon Imaging Center, Northwestern University). Serial optical sections (n=30-40; each slice is 0.1 mm thick) spaced by 0.2 mm were acquired. The data sets were deconvoluted using Imaris (version 9.0) software and optical sections were merged to produce 3D images. Spatial distances between probes were measured as previously described (Jhunjhunwala *et al.*, 2008; Montefiori *et al.*, 2016). Purified pro-B cells from at least two mice or two slides from cell lines of different genotypes were investigated and 302-404 alleles were analyzed. P values of statistical significance were calculated using Mann-Whitney U-test (Suppl. Table 7).

**RNA-seq**

Total RNA was prepared from Rag2$^{-/-}$ and Rag2$^{-/-}$NE1$^{-/-}$ pro-B cells. RNA was extracted using TRIzol (Life Technologies) and genomic DNA (gDNA) was eliminated using the genomic DNA wipeout buffer in the QuantiTect Reverse Transcription Kit (Qiagen). RNA samples were submitted to the Genome Research Core, UIC where RNA was purified using with the RNeasy Kit (Qiagen), ribosomal RNA was eliminated using the Ribo-Zero Magnetic Gold Kit (Illumina), processed with the NEBNext Ultra Directional RNA Library Prep Kit (Illumina) and sequenced on the Illumina HiSeq system (University of Illinois, Urbana). Three independent RNA-seq samples were prepared for Rag2$^{-/-}$ and Rag2$^{-/-}$NE1$^{-/-}$ pro-B cells. RNA-seq data have been deposited in the Gene Expression Omnibus (GEO) database (accession number GSE203402).

**V(D)J-seq**

Total BM cells were harvested from long bones of 4-5 mice that were 6 to 10 weeks old. CD19+ cells were isolated using anti-CD19 conjugated MACS beads (Miltenyi, Auburn CA) and then pre-incubated with CD16/32 Fc Block for 5 min and stained with antibodies for CD19-BB515, CD93-PE/Cy7, IgM Fab-AF647, CD2-PE, CD43-BV421 (Suppl. Table 1). Genomic DNA (gDNA) was isolated from purified pro-B cells (DNeasy, Qiagen) and the V(D)J-seq protocol performed while omitting the negative depletion step and using custom $J_H$ primers (Suppl. Table 8) (Bolland *et al.*, 2016; Matheson et al., 2017). GDNA was sonicated to a range of 500 bp to 1000 bp using a Covaris S2 (Covaris). Library barcoding was carried out using NEBNext Multiplex Oligos for Illumina (E7600S). Samples were paired-end 2x300 sequenced on an Illumina MiSeq System (San Diego, CA) at the NGS Core, Scripps Research. $V_H$ gene usage was determined using MIXCR software and data have been deposited in the GEO database (accession number: GSE203484).

**Statistics**

P values were calculated by using two-tailed Student's t test, Mann-Whitney U test or Mann-Whitney Wilcoxon test as indicated. In all cases $p>0.05$ (*), $p>0.01$ (**), $p>0.001$ (***) $p>0.0001$ (****).

**3C library construction and analysis**

3C chromatin was prepared from $CD19^{+}$ IL7 expanded $Rag2^{-/-}$ pro-B cells, the Abl-t 445.3.11 line and ConA activated splenic T cells. Optimized 3C library construction and assays for the Igh locus using Hind III were performed as described (Feldman et al., 2015; Feldman *et al.*, 2017). Briefly, Rag2-/- pro-B cell, the Abelson transformed (Abl-t) pro-B cell line 445.3.11 and ConA activated T cells were crosslinked using 1% formaldehyde and template concentration using the Mb1 primers was determined (Wuerffel et al., 2007). Quantitative PCR (qPCR) in combination with 5'FAM and 3'BHQ1 modified probes (IDT) was used to detect of 3C products and primers were designed using Primer Express software (ABI) (Suppl. Table 9). Primer and probe optimization were carried out according to the manufacturer's recommendations, (http://www3.appliedbiosystems.com/cms/groups/mcb_support/documents/generaldocuments/cms_042996.pdf). A template in which all possible 3C ligation products are present in equimolar concentration was used to control for differences in amplification efficiency between primer sets (Suppl. Table 9). The data were normalized using the interaction frequency between two fragments within the non-expressed Chr5 gene desert to facilitate sample to sample comparisons (Suppl. Table 9). The relative crosslinking frequency between two *Igh* restriction fragments was calculated: $X_{Igh} = [S_{Igh}/S_{GD}]$ Cell Type/$[S_{Igh}/S_{GD}]$ Control mix. $S_{Igh}$ is the signal obtained using primer pairs for two different *Igh* restriction fragments and $S_{GD}$ is the signal obtained with primer pairs for the *GD* locus fragments. The crosslinking frequency for the *GD* fragments was set to 1 to allow sample to sample comparisons. Data are represented as mean ± SEM. A complete laboratory protocol for 3C is available upon request. For all qPCR 3C reactions, 100 ng of chromatin was used.

**Hi-C library construction and analyses**

Genome-wide in situ HiC libraries were constructed from Rag1-/-, Rag1-/-NE1-/- pro-B cells expanded in IL7 for 4-5 days using Arima Hi-C kits (Arima Genomics, San Diego, CA) as recommended by the manufacturer. *In situ* Hi-C was performed using two biological replicates that yielded a minimum of 1.3 billion read pairs and 0.72 billion from pro-B cells of each genotype (Suppl. Table 10) (GEO Accession No. GSE201357). Published in situ Hi-C data sets for mouse embryonic fibroblasts (MEFs) were constructed with Arima Hi-C kits (GEO Accession No. GSE113339) (Di Giammartino *et al.*, 2019) and data was handled in parallel with the pro-B cell data. Hi-C data have been deposited in the GEO database (accession number GSE201357).

*In situ* Hi-C data was processed using the Juicer pipeline (v.1.5), CPU version (Durand et al., 2016). The pipeline was set up with BWA (0.7.15- r1140) (Li and Durbin, 2010) to map each read end separately to the mm10 reference genome (GRCm38). Duplicate and near-duplicate reads, as well as reads that map to the same fragment were removed. Among the remaining reads, those with mapping quality score (MAPQ) < 30 are retained. Arima-HiC-specific restriction sites (mm10_GATC_GANTC.txt) were obtained from ftp://ftp-arimagenomics.sdsc.edu/pub/JUICER CUTSITE FILES. For individual replicates, raw reads from each replicate were mapped separately to the reference genome, and then filtered by the MAPQ score. For merged replicates of each sample type, valid read pairs from both replicates were merged, mapped to the reference genome, and then filtered by the MAPQ score. Hi-C contact matrices were extracted from .hic files using the Dump command provided by a Java-based program in Juicer tools (Durand *et al.*, 2016). Hi-C maps were then normalized with the Knight and Ruiz (KR) matrix balancing method (Knight and Ruiz, 2013). Heatmap resolution of all individual replicates was computed using the Juicer script (calculate_map_resolution.sh) and found to be >10 kb. The reproducibility of Hi-C data was computed using the Stratum-adjusted correlation coefficient (SCC) on Chr12 using HiCRep at different resolutions (Yang et al., 2017). The replicates were merged and displayed at 10Kb resolution.

Extraction of virtual 4C interaction matrices: FASTQ files were converted to hic files using the pre-juicer commands from Juicebox (Dudchenko, 2018; Durand *et al.*, 2016). The hic files were used to generate virtual 4C viewpoints from dumped matrices generated in Juicebox. KR normalized observed read matrices were extracted at 10kb resolution. The biological replicates had stratum adjusted correlation coefficient (SCC) (Yang *et al.*, 2017) greater than 0.9 and were merged. The interaction profile of virtual 4C were plotted by running a rolling window of 30kb with a 10kb slide.

Generation of Hi-C difference maps: Experimentally measured Hi-C contact matrices of individual replicate and merged samples were quantile normalized against the Hi-C contact matrices of the uniformly sampled random ensemble of the corresponding cell type. Normalizing each sample against the target distribution of uniformly sampled random ensemble will remove the between-replicate biases in frequencies due to difference in sequencing depth (Dixon et al., 2015). These quantile normalized frequencies of each sample are then compared with each other, from which differential plots are obtained to identify differences between the two biological samples. Specifically, quantile normalized frequencies of MEF are subtracted from quantile normalized frequencies of each pro-B cell KO sample.

#### Supplementary Tables

**Supplementary Table 1:** Antibodies and reagents for analysis and isolation of pro-B and B cells.

| Antibodies /Reagents | Name/Clone | Source | Identifier |
|---|---|---|---|
| CD19-BB515 | BB515 Rat anti mouse CD19/1D3 | BD Biosciences | Cat# 564509; RRID:AB_11153299 |
| CD93 PECy7 | PE/Cyanine7 anti-mouse CD93/AA4.1 (RUO) | Biolegend | Cat#136506; RRID_AB_2044012 |
| IgM Fab-AF647 | Alexa Fluor® 647 AffiniPure Fab Fragment Goat Anti-Mouse IgM/ Polyclonal | Jackson Immuno Research | Cat#115-607-020; RRID:AB_2338932 |
| CD2-PE | PE anti-mouse CD2 antibody/RM2-5 (RUO) | Biolegend | Cat#100108; RRID:AB_2073690 |
| CD43-BV421 | BV421 Rat anti mouse CD43/S7 (RUO) | BD Biosciences | Cat # 562958; RRID:AB_2738069 |
| B220-APC-Cy7 | CD45R (B220) monoclonal antibody/RA3-6B2 (APC-eFluor 780) | eBioscience | Cat # 47-0452-82 RRID:AB_1518810 |
| CD19-PerCp | PerCP anti-mouse CD19 antibody/6D5 (RUO) | Biolegend | Cat # 115532 RRID:AB_2072926 |
| CD43 (S7)-APC | APC rat anti-mouse CD43/S7 (RUO) | BD Biosciences | Cat # 560663 RRID:AB_1727479 |
| IgM-e450 | IgM Monoclonal antibody/ eB121-15F9 (eFluor 450) | eBioscience™ | Cat # 48-5890-82 RRID:AB_10671539 |
| CD93-PE Cy7 | CD93 monoclonal antibody/AA4.1 (PE-Cyanine7) | eBioscience™ | Cat # 25-5892-81 RRID:AB_469658 |
| IgD-AF700 | Alexa Fluor® 700 anti-mouse IgD antibody/11-26c.2a (RUO) | Biolegend | Cat # 405730 RRID:AB_2563341 |
| CD5-PE | PE anti-mouse CD5 antibody/53-7.3 (RUO) | Biolegend | Cat # 100608 RRID:AB_312737 |
| CD23-biotin | Biotin Rat Anti-Mouse CD23 /B3B4 (RUO) | BD Biosciences | Cat #553137 RRID:AB_394652 |
| CD21-FITC | Rat Anti-CD21 / CD35 Monoclonal Antibody/7G6 | BD Biosciences | Cat #553818 RRID:AB_395070 |
| Streptavidin-APC | Streptavidin-allophycocyanin antibody | BD Biosciences | Cat #554067 RRID:AB_10050396 |
| CTCF | Rabbit Anti-CTCF Polyclonal antibody | MilliporeSigma | Cat# 07-729 RRID:AB_441965 |
| CD19 conjugated MACS beads | CD19 microbeads mouse | Miltenyi Biotech | Cat# 130-052-201 |
| FVS-510* | Fixable Viability Stain 510 (RUO) | BD Horizon™ | Cat # 564406 RRID:AB_2869572 |
| CD16/32 Fc Block | Purified anti-mouse CD16/32 Antibody/93 (RUO) | Biolegend | Cat# 101302; RRID:AB_312801 |

**Supplementary Table 2:** Primers used for Igh gene expression qPCR and qRT-PCR assays.

| Name | Sequence | Reference |
|---|---|---|
| Iγ2b F | ATCCCAGAGTCACAGAGGAA | (Shen et al., 2021) |
| Cγ2b R | CACACCTACAGACAACCAGAC | (Dai *et al.*, 2021) |
| Iµ F | TCCACACAAAGACTCTGGACC | (Shen *et al.*, 2021) |
| Cµ R | TCAGTGTTGTTCTGGTAGTTCCAG | (Shen *et al.*, 2021) |
| µ0µ F | GAAGACATTTGGGAAGGACTGA | This study |
| µ0Cµ R | GAACAGAGGCAGAACAGAGAC | This study |
| $V_H$14-2 F | TGGAATTTCGGGGCATATTTAGTTTCACC | This study |
| $V_H$14-2 R | TCATCTTCTTCCTGATGGCAGTGGTTAC | This study |
| DQ52 F | CGGACAGAGCAGGCAGGTGG | (Oudinet *et al.*, 2020) |
| DQ52 R | GCATCCAAGCCTCAGAAACTCAG | (Oudinet *et al.*, 2020) |
| $V_H$81X F | AAGTGGGGGACGAGGAAGAC | (Oudinet *et al.*, 2020) |
| $V_H$81X R | CTGGGGGGGTGTGTTTCC | (Oudinet *et al.*, 2020) |
| 7183 F | CACAGTGAGATTCAGAACACCCTTA | (Oudinet *et al.*, 2020) |
| 7183 R | GAAATGAGGAAGGCAGGCG | (Oudinet *et al.*, 2020) |
| J606 F | AGGTTAGTCTGGTGAGGCATA | (Braikia et al., 2015) |
| J606 R | CCAACCTACTCTAACCTCTGCTA | (Braikia *et al.*, 2015) |
| J558 F | ATTCCCCTCCCAATAGGAAA | (Bolland *et al.*, 2004) |
| J558 R | TGTCAATCACAATGGGCATC | (Bolland *et al.*, 2004) |
| VHJ558 GLT F | ATGGGATGGAGCTGGATCTT | (Guo et al., 2011) |
| VHJ558 GLT R | CTCAGGATGTGGTTACAACACTGTG | (Perlot et al., 2005) |
| PAIR4 F | ATGGGGCACATAGGTTCTTCC | (Puget et al., 2015) |
| PAIR4 R | GGACATCTGAGAGATCATTGAACATC | (Puget *et al.*, 2015) |
| VH81XRT FP | CCCAAGACATGTCATGGGAAGGGAATTC | This study |
| VH81XRT RP | ATGGACTTCGGGCTCAGCTTGG | This study |
| VHQ52.7.18 RT FP | CAGTCTGGACATGAAAGCTGCATTG | This study |
| VHQ52.7.18 RT RP | CTGTCCTGGTGCTGCTCC | This study |
| 18SF | TTGACGGAAGGGCACCACCAG | (Rhinn et al., 2008) |
| 18SR | GCACCACCACCCACGGAATCG | (Rhinn *et al.*, 2008) |

**Supplementary Table 3:** Guide RNAs for CRISPR/Cas9 genome editing.

| Sites | gRNA | Sequence (Bold = PAM) | Genomic Coordinates (mm9) | Clones/Mice |
|---|---|---|---|---|
| SiteI.3 | Site I.3 g5 | TTACATAAGAACATTCTGGC**TGG** | | $V_H$14-2 Pr KO.1, $V_H$14-2 Pr KO.2 |
| | Site I.3 g6 | GACTGTGATGATTAATATAT**AGG** | | |
| F.6 | F.6 g1 | CTAAGATGCTCATGGTCAGC**AGG** | | F.6 KO.1, F.6 KO.3 |
| F.6 | F.6 g2 | TGACCATGAGCATCTTAGAG**TGG** | | F.6 KO.2 |
| | F.6 g3 | GCTGGAACATAAGGAGCTCC**AGG** | | |
| NE 1 | NE1 g1 | GGGGAAATTGCATTGTAACA**TGG** | | NE1 KO.1, NE1 KO.2 |
| | NE1 g3 | GGGTTATTCTTGTGTGACCC**AGG** | | NE1 KO.3 |
| NE2 | NE2 g1 | GTCAGACCCCTTAACTGACC**AGG** | | NE2 KO.1, NE2 KO.2 |
| | NE2 g2 | AAGTGTTGAAGTAGTGCAC**AGG** | | NE2 KO.3, NE2 KO.4 |
| NE1 | NE1 g1 | GGGGAAATTGCATTGTAACA**TGG** | | NE 1 $^{-/-}$ mice |
| | NE1 g3 | GGGTTATTCTTGTGTGACCC**AGG** | | |

**Supplementary Table 4:** Primers used for genotyping CRISPR/Cas9 edited cell lines and mice

| Name | Sequence | Reference |
|---|---|---|
| c_SiteI.3 F1 | AGTGAAGAGCTGGGCAAAATATAGC | This Study |
| c_SiteI.3 R1 | GAATGCACATATTTCTGAGGCATG | This Study |
| c_SiteI.3 F2 | ATTTGCATACTCATGAGGCAGGATC | This Study |
| c_SiteI.3 R2 | CTGCACATGTCTCAGCGAAAT | This Study |
| c_F.6 F | TTCTGTCTGAGCTCCCAATTTACTTTCCTT | This Study |
| c_F.6 R | GGAGAAGGATTGTGAATTGTAAAATTACTCTAAATGGTT | This Study |
| c_NE1 F1 | GTCTAAACATATTTTGTAGGGTA | This study |
| c_NE1 R1 | GATCACTTACTCTCAAATCGAGAC | This study |
| c_NE1 EP F | TTGTCTAAACATATTTTGTAGGGTATATTTGAA | This study |
| c_NE1 EP R | GGTTCATTGAGTCACAAGGTAGTTC | This study |
| c_NE1 IP F | CCATGTTACAATGCAATTTCCCC | This study |
| c_NE1 IP R | CTGGGTCACACAAGAATAACCC | This study |
| c_NE2 F | AAACATTTGAGGTACAGAACAT | This Study |
| c_NE2 R | TCTAGGAAGACCTATTATTGCT | This Study |
| Rag2 WT | TCGATTCCTAGAGCGTCCTT | The Jackson Laboratory |
| Rag2 KO | GGTCATCCTTTGCAACACAG | The Jackson Laboratory |
| Rag2 Common | CAGCGCTCCTCCTGATACTC | The Jackson Laboratory |
| Rag1 0189 | TGGATGTGGAATGTGTGCGAG | The Jackson Laboratory |
| Rag1 3104 | CCGGACAAGTTTTTCATCGT | The Jackson Laboratory |
| Rag1 1746C | GAGGTTCCGCTACGACTCTG | The Jackson Laboratory |

**Supplementary Table 5:** Primers used for $D_H$->$J_H$ and $V_H$->$D_HJ_H$ recombination assays.

| Name | TaqMan Probe | Sequence | Reference |
|---|---|---|---|
| VH81XF | | GAAACTCTCCTGTGAATCCAATGAATACGAA | This study |
| VH7183-VH81X F | | TGTGCAGCCTCTGGATTCACT | (Espinoza and Feeney, 2005) |
| VHJ558 | | CCTCCARCACAGCCTACATGSA | (Volpi et al., 2012) |
| VH7183 F | | CCGATTCACCATCTCCAGAGAC | (Volpi *et al.*, 2012) |
| DFL16.1 F | | ACACCTGCAAAACCAGAGACCATA | (Subrahmanyam et al., 2012) |
| JH1 R | | CCCAGACATCGAAGTACCAGTAG | This study |

**Supplementary Table 6:** Primers used for the short FISH probes

| **Name** | **Primer Sequence** | **Reference** |
|---|---|---|
| Eμ F | CCCACAGGCTCGAGAACTTTAGCGAC | (Gerasimova et al., 2015) |
| Eμ R | GCTGGAGAGTTAGTCCAGCCGAC | This study |
| Site I.3 F | GAGTAAAAAGGAATCACCCTTTTAAAACTAAGC | This study |
| Site I.3 R | TGAGTGTCCTTAAGTCTGCTGC | This study |
| NE1 F | TAGATCCCTGAAAGATGAGATTG | This study |
| NE1 R | CAGTTGAACCCAGAGGTAAG | This study |
| NE2 F | GCTCCAAGCAGAGAGTGTATTC | This study |
| NE2 R | CTCAGGCCCTCAAGTGTTATTC | This study |

**Supplementary Table 7. Statistical analysis of pairwise FISH probe contacts[1].**

| Experiments | Probes | Samples | Alleles measured[2] | P value[3] |
|---|---|---|---|---|
| | | **Mouse** | | |
| 1 | BAC H14 - RI | Rag2-/- vs Rag2-/- NE1-/- | (H14, RI) 400, 400 | 0.1702 |
| 2 | BAC RI - RII - RIII | Rag2-/- vs Rag2-/- NE1-/- | (RI, RII) 400, 400 | 0.0004 |
| | | | (RII, RIII) 400, 400 | 0.0890 |
| | | | (RI, RIII) 400, 400 | <0.00001 |
| 3 | E$\mu$ - NE1 - NE2 | Rag2-/- vs Rag2-/- NE1-/- | (E$\mu$, NE1) 400, 400 | 0.7231 |
| | | | (NE1, NE2) 400, 400 | 0.0312 |
| | | | (E$\mu$, NE2) 400, 400 | <0.00001 |
| | | **Cell lines** | | |
| 4.1 | BAC H14 - RI | 445.3.11 vs $V_H$ Pr KO | (H14, RI) 302, 302 | 0.9112 |
| 4.2 | BAC H14 - RI | 445.3.11 vs NE1 KO | (H14, RI) 302, 302 | 0.8833 |
| 4.3 | BAC H14 - RI | 445.3.11 vs F.6_CBE KO | (H14, RI) 302, 302 | 0.0513 |
| 5.1 | BAC RI – RII - RIII | 445.3.11 vs $V_H$ Pr KO | (RI, RII) 302, 302 | 0.2262 |
| | | | (RII, RIII) 302, 302 | <0.00001 |
| | | | (RI, RIII) 302, 302 | 0.0433 |
| 5.2 | BAC RI - RII - RIII | 445.3.11 vs NE1 KO | (RI, RII) 302, 302 | <0.00001 |
| | | | (RII, RIII) 302, 302 | <0.00001 |
| | | | (RI, RIII) 302, 302 | <0.00001 |
| 5.3 | BAC RI - RII - RIII | 445.3.11 vs F.6_CBE KO | (RI, RII) 302, 302 | 0.0002 |
| | | | (RII, RIII) 302, 302 | 0.0057 |
| | | | (RI, RIII) 302, 302 | <0.00001 |
| 6.1 | E$\mu$ - Site I.3 - NE1 | 445.3.11 vs $V_H$ Pr KO | (E$\mu$, Site I.3) 404, 404 | 0.0015 |
| | | | (Site I.3, NE1) 404, 404 | <0.00001 |
| | | | (E$\mu$, NE1) 404, 404 | 0.0049 |
| 6.2 | E$\mu$ - Site I.3 - NE1 | 445.3.11 vs NE1 KO | (E$\mu$, Site I.3) 404, 404 | <0.00001 |
| | | | (Site I.3, NE1) 404, 404 | <0.00001 |
| | | | (E$\mu$, NE1) 404, 404 | 0.0423 |
| 6.3 | E$\mu$ - Site I.3 - NE1 | 445.3.11 vs F.6_CBE KO | (E$\mu$, Site I.3) 404, 404 | 0.0088 |
| | | | (Site I.3, NE1) 404, 404 | <0.00001 |
| | | | (E$\mu$, NE1) 404, 404 | <0.00001 |
| | | **Mouse vs Cell line** | | |
| 7 | BAC H14 - RI | Rag2-/- vs 445.3.11 | (H14, RI) 400, 302 | <0.00001 |
| 8 | BAC RI – RII - RIII | Rag2-/- vs 445.3.11 | (RI, RII) 400, 302 | <0.00001 |
| | | | (RII, RIII) 400, 302 | 0.00018 |
| | | | (RI, RIII) 400, 302 | 0.00016 |

**1** Related to Figures 3, 6, 7 and Figure SX.

**2** The number of alleles for each genotype and probe pair.

**3** P-values are reported comparing the spatial distance between two probes and are calculated with the Mann-Whitney U-test using GraphPad Prism software.

**Supplementary Table 8:** Primers for VDJ-seq.

| **Oligonucleotides** | **Sequence** | **Reference** |
|---|---|---|
| **Adapter** | | |
| VDJseq F (Mix 1) | ACACTCTTTCCCTACACGACGCTCTTCCGATCTNNNNNNGACTCG*T | (Matheson *et al.*, 2017) |
| VDJseq R (Mix 1) | /5Phos/CGAGTCNNNNNNAGATCGGAAGAG*C/3SpC3/ | (Matheson *et al.*, 2017) |
| VDJseq F (Mix 2) | ACACTCTTTCCCTACACGACGCTCTTCCGATCTNNNNNNCTGCTCC*T | (Matheson *et al.*, 2017) |
| VDJseq R (Mix 2) | /5Phos/GGAGCAGNNNNNNAGATCGGAAGAG*C/3SpC3/ | (Matheson *et al.*, 2017) |
| **Enrichment** | **Biotinylated extension primer sequence** | |
| JH1 enrichment R | /5Biosg/AGCCAGCTTACCTGAGGAGAC | This study |
| JH2 enrichment R | /5Biosg/GAGAGGTTGTAAGGACTCACCTG | This study |
| JH3 enrichment R | /5Biosg/AGTTAGGACTCACCTGCAGAGAC | This study |
| JH4 enrichment R | /5Biosg/AGGCCATTCTTACCTGAGGAG | This study |
| **PCR Primers** | | |
| Short PE1/i5 | ACACTCTTTCCCTACACGACGCTC*T | (Kleiman et al., 2018) |
| $J_H$1/i7 at 5' end | GTGACTGGAGTTCAGACGTGTGCTCTTCCGATCT TTACCTGAGGAGACGGTGACC*G | This study |
| $J_H$2/i7 at 5' end | GTGACTGGAGTTCAGACGTGTGCTCTTCCGATCT GGACTCACCTGAGGAGACTGTG*A | This study |
| $J_H$3/i7 at 5' end | GTGACTGGAGTTCAGACGTGTGCTCTTCCGATCT GGACTCACCTGCAGAGACAGTGA*C | This study |
| $J_H$4/i7 at 5' end | GTGACTGGAGTTCAGACGTGTGCTCTTCCGATCT CCATTCTTACCTGAGGAGACGGTG*A | This study |

**Supplementary Table 9:** Primers and probes used for 3C assays.

| Primer | TagMan Probe | Sequence | Reference |
|---|---|---|---|
| E*μ* | | TCCACACAAAGACTCTGGACCTCT | (Feldman *et al.*, 2015) |
| | **A (E*μ*)** | TGGCTTACCATTTGCGGTGCCTGGTTT | (Feldman *et al.*, 2015) |
| T.H (hs3b,4) | | GCCCCTAAGACCCTACTCTGCTA | (Feldman *et al.*, 2015) |
| | **H (hs3b,4)** | TGACTCATCCACATCACCTTGCCT | (Feldman *et al.*, 2015) |
| IGCR1 | | AGCCTCAAGTTCCTTCAGGGC | This study |
| VH81X | | GAACATTCTGGGAACAGTATTAGGATG | This study |
| Q52.2.4 | | CTGGATGCCTGTCTCCTGTAG | This study |
| Q52.3.8 | | CTACTGTGGTAGCTTCTACCCG | This study |
| 7183.7.10 | | GTGAAGGACTTTGCAGGTTGGAC | This study |
| Q52.5.13 | | AAGACTCCCATTCTCTGAGACTGG | This study |
| 7183.14.25 | | TTGCACTGCACATCACTGCTGG | This study |
| Ia1 | | GGAAGGGAATGGGCATTAATGCATTAAG | This study |
| Ia2 | | CAGCTGTAGTGATGGTAATAAAGTG | This study |
| Ia3 | | CGAGTAAATGGGGACAGGGGAAAT | This study |
| Ia4 | | CTTAGCCTTTCGGTATTCCTCC | This study |
| Ia5 | | GGTATCCAGAACTTGATGTGGTGA | This study |
| I.1 | | GTCGTCATCTACACACAGGAGC | This study |
| I.1a | | GAAAACAAACTCAAAGACAAACTACATGATA | This study |
| I.1b | | CCAGATACATAGAATAGGAGAGAAGAC | This study |
| I.1c | | CGGAGAAGATCAAATTGAAAATCCTCTC | This study |
| I.1d | | GGACCCATCCAACCATCTTTGAAATG | This study |
| I.1e | | GACTGTTGTTGACAGATTTGGCATTC | This study |
| I.1f | | CTGGACTGTAAAGCCAGTTCCAC | This study |
| I.1g | | CTGGAGTCTGGCACCATCATCAT | This study |
| I.1h | | GATAAAACTGAAGGTCTGCCCACTC | This study |
| I.1i | | CTCTCTGAGCTCAGGAATTCCAG | This study |
| I.1j | | CAAGAACAAATTTGAGTCAGATCTTCTG | This study |
| I.2 | | CGTTATGCTTTTAATTTGCTCACTTATAAG | This study |
| I.2a | | CAATACCCATATGGTATTTTCAACAATTTTTG | This study |
| I.3 | | TCCTACTTCTGTCTGAGGAAATTTCG | This study |
| I.3a | | CTCAGGAATTCGAGCAGAAGAT | This study |
| I.3b | | ACAAACTCTGAAGAAGGTCCGAAGGT | This study |
| | **P_Site I.1** | CCTGTGGGACTCTAAGCCCATGTCAG | This study |
| | **P_Site I.3** | TGATATTGTATCTAAGTGGTCTGCACATGTC TCAG | This study |
| 5'F.0 | | ATTCTTCTCAATCCTGGGACTAGT | This study |
| F.0 | | TGTCTGCATGGTGTTTCTTTCATAC | This study |
| F.1 | | TAGTGCAGCTTCCACTTAAAGAAC | This study |
| F.2 | | GACACCATGGAGATGGATCATC | This study |
| F.3 | | CCTCCTGCAGCAGGGTTATTC | This study |
| F.4 | | CCCACAAATGCCAGACTAAAGAAC | This study |
| F.5 | | CACTTTGAAGCTGAAGACATTGAC | This study |
| F.6 | | CTGAAAGCAGGCTATTCACCATG | This study |
| F.7 | | GCAAAGGACCTCTAAAGGATTGC | This study |
| F.8 | | GATTCCATAAACAGAACCCTAATGG | This study |

| | | | |
|---|---|---|---|
| F.9 | | GCCTGTTTGTTGAGAAGTGCACA | This study |
| | **P_F.6** | AGAGCAACAAGGAAAAGCCATCTAAGCTCC AA | This study |
| Fb.1 | | TTACCCACTCTATCCAGTAGCC | This study |
| Fb.2 | | CGTTCAGAGCAACACTGCCCTA | This study |
| Fb.3 | | ACTTGGAGACATCTAAGAAAAGTAAGATTAG | This study |
| Fb.4 | | CTTAGAGGACTACTTTCATAATTGAATTCC | This study |
| Fb.5 | | ATCCCTACTCTTGAATCTCAATGAGG | This study |
| Fb.6 | | ATTTGCAGAAAGTTGAGTAGAAAGAACAG | This study |
| Fb.7 | | CGTTTGTGCTATATGATAGAGATTTGCCA | This study |
| Fb.8 | | CTGCTAGAGTAGCCCTTGTCCATAA | This study |
| Fb.9 | | GTGAACATTACCACATGAGGGACTTC | This study |
| IIa1 | | CATCAAGTATCAATAAAAGAGACCTCATAAA G | This study |
| IIa2 | | GAAACGCCAATCTGACTTCCAGTG | This study |
| IIa3 | | AGAGAAACACTCCTCCACTTCCG | This study |
| IIa | | GATGTGTAACTGTCTCCGTAATAATATATAG | This study |
| IIa.b | | CATCCCAGACAGAGCTACAGAG | This study |
| IIa.c | | TGGGAGAAGTTGCACTTACATCTTG | This study |
| IIa.d | | ACTCTTATATTCTAAAAACCTGGAACTTG | This study |
| IIa.e | | TCAAAGCAACCCTAAGAATCTCCTTC | This study |
| | **P_Site IIa** | TCTTCCCTCCCAAGCATGTACTCA | This study |
| | **P_Site IIa.b** | ACTCTGTAGCTCTGTCTGGGATGCT | This study |
| GD F | | AGGCTTCTGACCTGCATCTTGA | (Kumar *et al.*, 2013) |
| GD R | | TTCCAGAGCATTGTCAGCAAA | (Kumar *et al.*, 2013) |
| | **GD** | ACCTTGCTACTCTTCCCTGGTGTTTGTTGG | (Kumar *et al.*, 2013) |
| Mb1 F | | CCACGCACTAGAGAGAGACTCAA | (Wang et al., 2006) |
| Mb1 R | | CCGCCTCACTTCCTGTTCAGCCG | (Wang *et al.*, 2006) |
| C14 F (hs3b,4) | | ATAGGCCCTCCTCCCACCA | (Feldman *et al.*, 2015) |
| C14 R (hs3b,4) | | CATGCGTTTGTGTCTACATGC | (Feldman *et al.*, 2015) |
| C3 F (Eu) | | GGTATCAAAGGACAGTGCTTAG | This study |
| C3 R (Eu) | | CAGCCTAGTTTAGCTTAGCG | This study |

#### Suppl. Table 10. In situ Hi-C Library Statistics

| | Rag1-/-1 | Rag1-/-2 | Rag1-/-_merged_replicates |
|---|---|---|---|
| Sequenced Read Pairs | 704,893,796 | 595,148,192 | 1,300,041,988 |
| Normal Paired | 145,215,091 (20.60%) | 147,945,053 (24.86%) | 293,160,144 (22.55%) |
| Chimeric Paired | 475,045,221 (67.39%) | 376,176,420 (63.21%) | 851,221,641 (65.48%) |
| Chimeric Ambiguous | 81,395,262 (11.55%) | 68,035,698 (11.43%) | 149,430,960 (11.49%) |
| Unmapped | 3,238,222 (0.46%) | 2,991,021 (0.50%) | 6,229,243 (0.48%) |
| Alignable (Normal+Chimeric Paired) | 620,260,312 (87.99%) | 524,121,473 (88.07%) | 1,144,381,785 (88.03%) |
| Unique Reads | 478,530,695 (67.89%) | 414,774,186 (69.69%) | 885,121,908 (68.08%) |
| PCR Duplicates | 136,259,276 (19.33%) | 104,319,997 (17.53%) | 248,853,403 (19.14%) |
| Optical Duplicates | 5,470,341 (0.78%) | 5,027,290 (0.84%) | 10,406,474 (0.80%) |
| Library Complexity Estimate | 1,173,386,201 | 1,111,955,039 | 2,189,931,487 |
| Below MAPQ Threshold | 85,367,316 (12.11% / 17.84%) | 75,979,254 (12.77% / 18.32%) | 160,175,973 (12.32% / 18.10%) |
| Hi-C Contacts | 393,163,379 (55.78% / 82.16%) | 338,794,932 (56.93% / 81.68%) | 724,945,935 (55.76% / 81.90%) |
| Inter-chromosomal | 60,362,190 (8.56% / 12.61%) | 53,686,252 (9.02% / 12.94%) | 114,046,736 (8.77% / 12.88%) |
| Intra-chromosomal | 332,801,189 (47.21% / 69.55%) | 285,108,680 (47.91% / 68.74%) | 610,899,199 (46.99% / 69.02%) |
| Short Range (<20Kb) | 124,066,290 (17.60% / 25.93%) | 99,751,406 (16.76% / 24.05%) | 216,820,141 (16.68% / 24.50%) |
| Long Range (>20Kb) | 208,725,177 (29.61% / 43.62%) | 185,347,443 (31.14% / 44.69%) | 394,059,576 (30.31% / 44.52%) |

| | Rag1-/-NE1-/-1 | Rag1-/-NE1-/-2 | Rag1-/-NE1-/-_merged_replicates |
|---|---|---|---|
| Sequenced Read Pairs | 761,741,401 | 736,789,530 | 1,498,530,931 |
| Normal Paired | 170,981,180 (22.45%) | 161,579,433 (21.93%) | 332,560,613 (22.19%) |
| Chimeric Paired | 501,909,501 (65.89%) | 490,577,891 (66.58%) | 992,487,392 (66.23%) |
| Chimeric Ambiguous | 85,210,925 (11.19%) | 81,411,068 (11.05%) | 166,621,993 (11.12%) |
| Unmapped | 3,639,795 (0.48%) | 3,221,138 (0.44%) | 6,860,933 (0.46%) |
| Alignable (Normal+Chimeric Paired) | 672,890,681 (88.34%) | 652,157,324 (88.51%) | 1,325,048,005 (88.42%) |
| Unique Reads | 517,825,831 (67.98%) | 507,477,961 (68.88%) | 1,014,856,941 (67.72%) |
| PCR Duplicates | 148,381,644 (19.48%) | 138,830,218 (18.84%) | 297,781,998 (19.87%) |
| Optical Duplicates | 6,683,206 (0.88%) | 5,849,145 (0.79%) | 12,409,066 (0.83%) |
| Library Complexity Estimate | 1,264,111,007 | 1,280,223,735 | 2,436,635,481 |
| Below MAPQ Threshold | 92,053,634 (12.08% / 17.78%) | 89,227,706 (12.11% / 17.58%) | 179,928,939 (12.01% / 17.73%) |
| Hi-C Contacts | 425,772,197 (55.89% / 82.22%) | 418,250,255 (56.77% / 82.42%) | 834,928,002 (55.72% / 82.27%) |
| Inter-chromosomal | 42,925,248 (5.64% / 8.29%) | 42,718,155 (5.80% / 8.42%) | 85,641,452 (5.72% / 8.44%) |
| Intra-chromosomal | 382,846,949 (50.26% / 73.93%) | 375,532,100 (50.97% / 74.00%) | 749,286,550 (50.00% / 73.83%) |
| Short Range (<20Kb) | 137,296,147 (18.02% / 26.51%) | 135,640,527 (18.41% / 26.73%) | 263,861,777 (17.61% / 26.00%) |
| Long Range (>20Kb) | 245,542,117 (32.23% / 47.42%) | 239,883,125 (32.56% / 47.27%) | 485,407,716 (32.39% / 47.83%) |

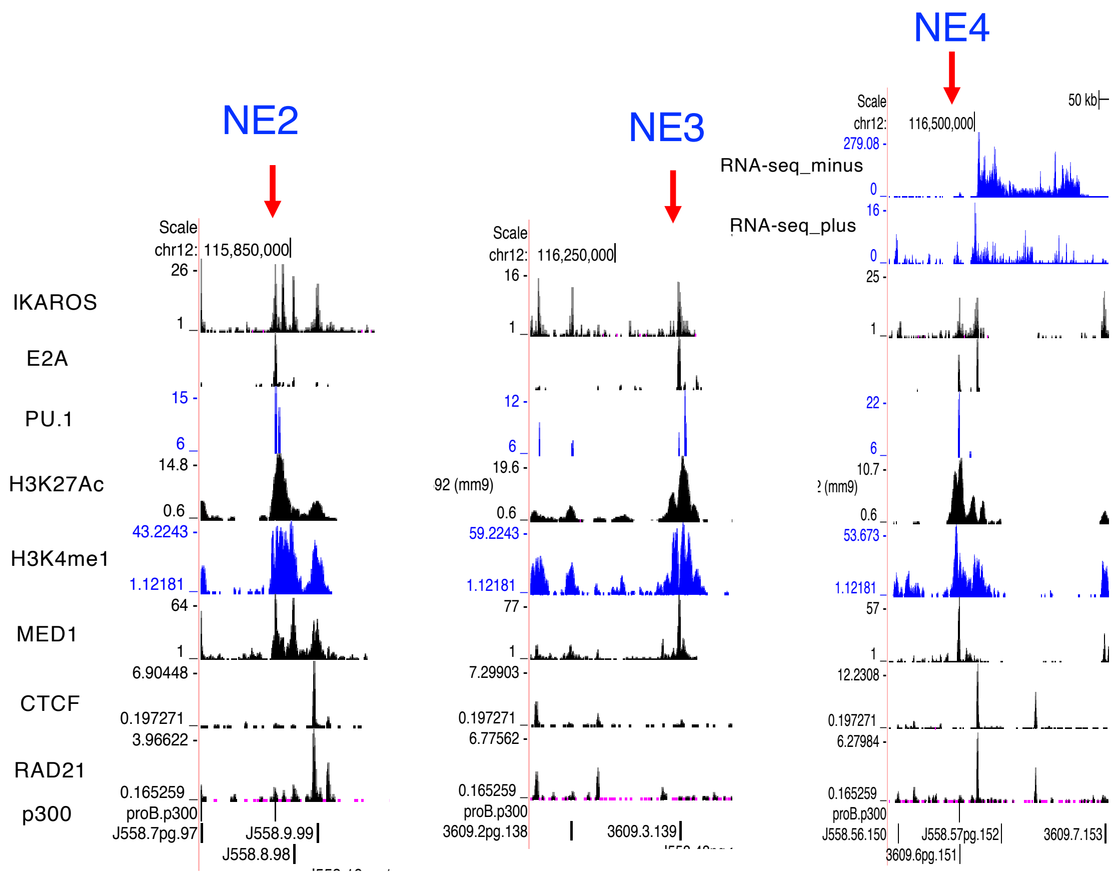


**Suppl. Figure 1. Histone protein modifications and protein factor binding signals for NE2, NE3 and NE4 in the Igh locus (related to Figure 1).** ChIP-seq data for histone modifications H3K27Ac, H3K4me1 and transcription factors p300, RAD21, CTCF, MED1, PU.1, E2A and IKAROS from Rag deficient pro-B cells are shown. RNA-seq data (+ and – strand) is shown for NE4. The data was compiled in UCSC genome browser using mm9 genome assembly.

### Suppl. fig. 1

A

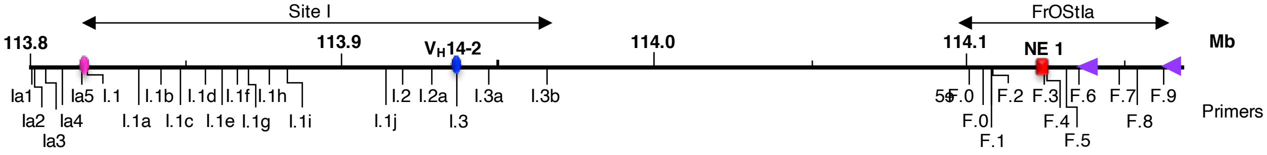


B

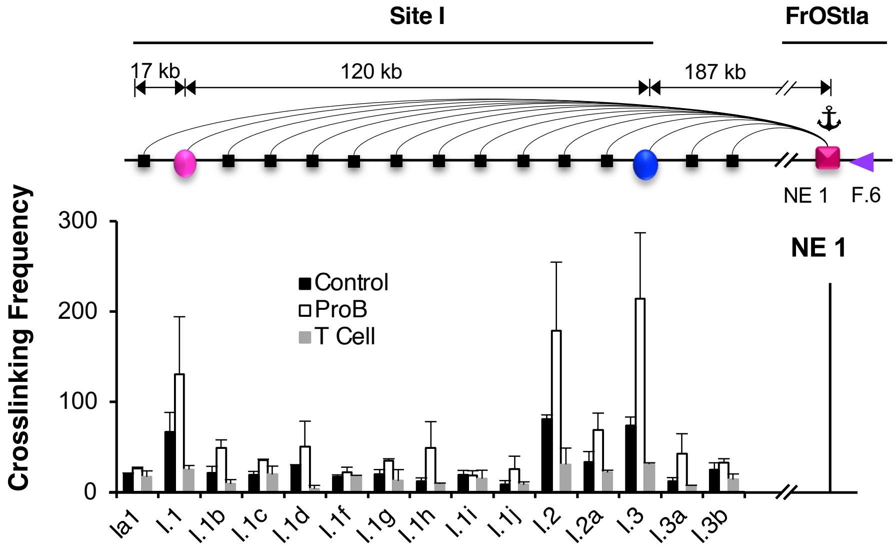


**Suppl. Figure 2. NE1 anchors chromatin loops with Site I (related to Figure 1). A)** Schematic of primer sites in Site I through FrOStIa highlighting DNA elements ( Site I.1, pink dot; Site I.3, blue dot; NE1, red square; F.6 CBE, purple arrow). Primers positions are labeled and represented by the vertical bars below the line. **B)** *Upper panel:* Arcs represent the 3C assays, primers are identified below the histograms, and anchor probes are indicated (anchor symbol). *Lower panels* 3C interaction profile using NE1 anchor probe and scanning the Site I region. Average crosslinking frequencies were derived from at least two independent chromatin preparations from Rag2$^{-/-}$ pro-B cells, ConA stimulated splenic T cells and Abelson transformed Rag1$^{-/-}$ 445.3 pro-B cell line and SEMs are shown. Two-tail Student's t test (*p<0.05; **p<0.001).

Suppl. fig 2

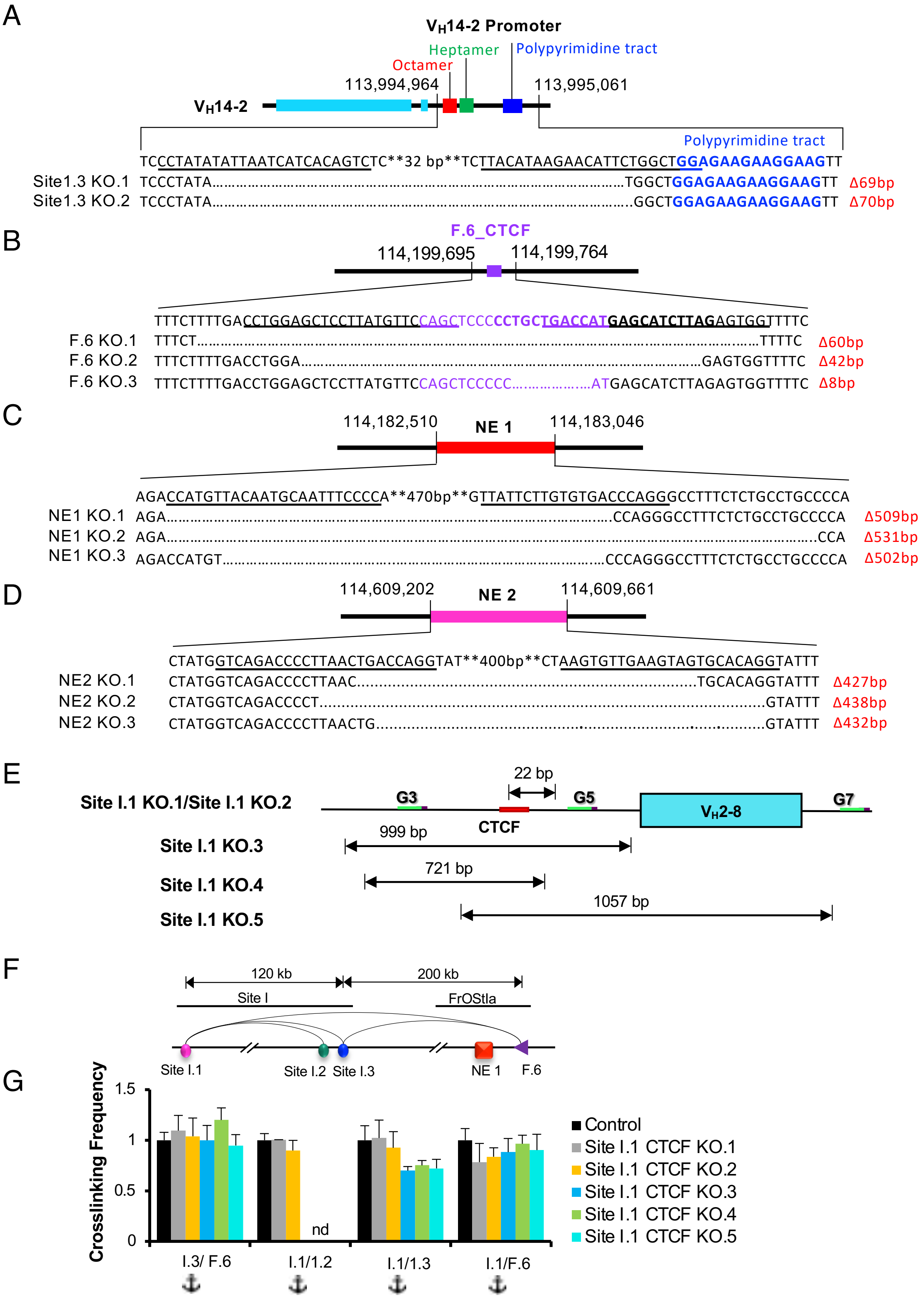


**Supplemental Fig. 3.**

**Suppl. Figure 3. CRISPR/Cas9 genome editing of TEs in a Abl-t pro-B cell line (related to Figure 2).** Genomic coordinates (chr12, mm9). **A-D)** Guide (g)RNA sequences are underlined. Deletions (Δ) and their positions are displayed below the parental DNA sequence. **A**) The $V_H$14-2 exon and leader (cyan rectangles), octamer (red rectangle), heptamer (green rectangle), polypyrimidine tract (blue rectangle), **B**) F.6 CBE (purple rectangle). KO.1 and KO.3 were obtained using a single gRNA indicated in bold. KO.2 clone was obtained using two guides. **C**) NE1 (red rectangle). **D**) NE2 (white rectangle). **E)** Summary of the CRISPR editing at Site I.1. Guide RNAs (G3, G5, G7) used for the editing, arrows indicate the positions of deletion, numbers on the arrows indicate the base pairs deleted, red horizontal bar shows CBE, teal rectangle indicates the $V_H$2-8 gene. **F)** Summary of the major loops tested in 3C assays (arcs) with genomic distances between them. **G)** Average crosslinking frequencies anchored at Site I.3 and Site I.1 from two independent chromatin templates prepared from control and KO clones. The data was normalized to 1 for controls. (nd, not done).

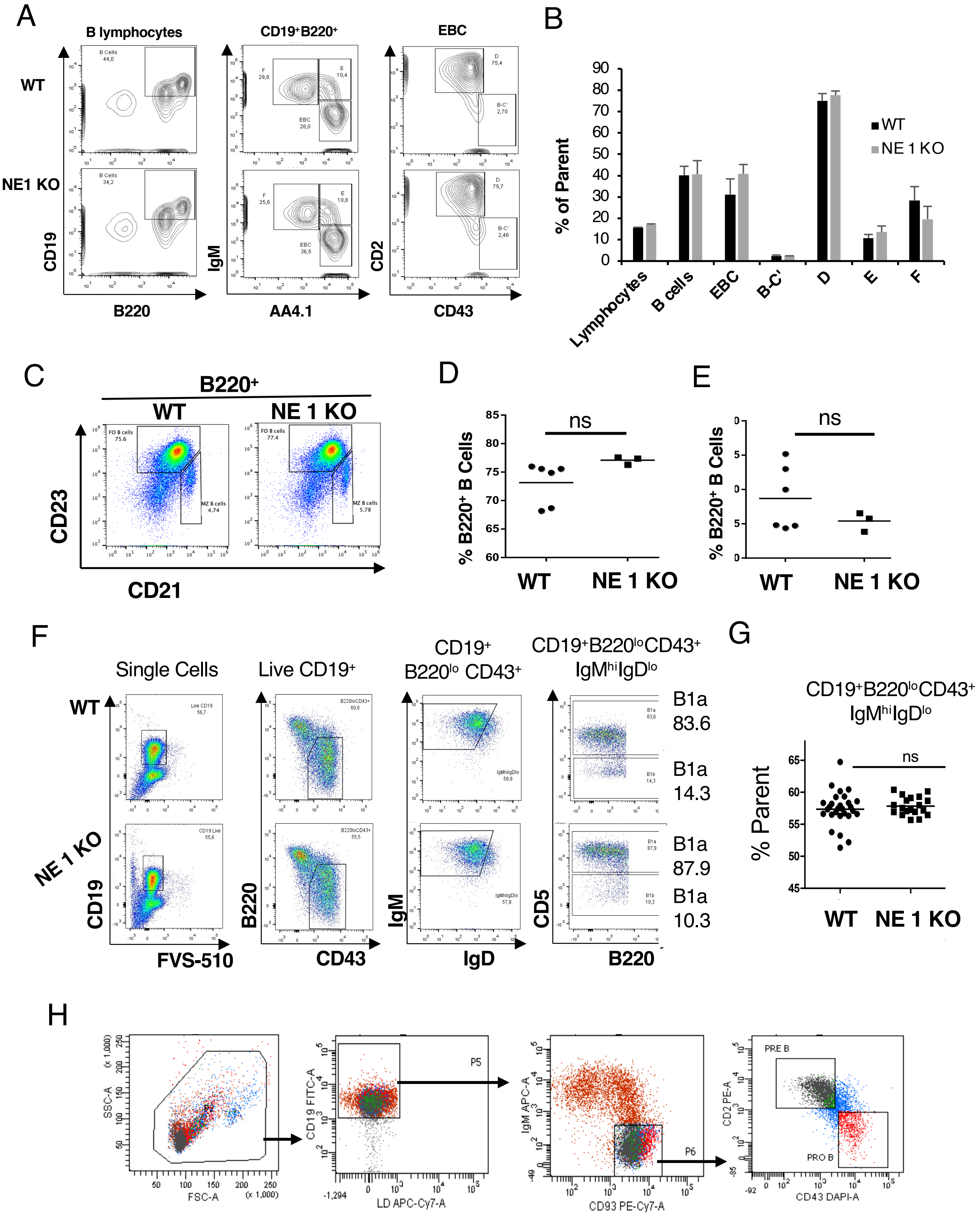


Suppl. Figure 4

**Suppl. Figure 4. B cell compartments are normal in NE1-/- mice. A) Analysis of Hardy fractions in BM (related to Figure 4).** BM was collected from humerus, tibia and femur bones of WT, and NE $1^{-/-}$ mice. Cells were stained with antibodies (CD19-PerCp, B220-APC Cy7, IgM-e450, AA4.1 PE Cy7, CD43 (S7)-APC, CD2-PE). Representative histograms depicting different B cell fractions. B lymphocytes (B220+CD19+), EBC (early B cells, B220+IgM-AA4.1$^{+}$ (CD93$^{+}$)), Hardy fractions B-C' (pro-B cells), D (pre-B cells), E (immature B cells) F (mature B cells). **B)** The graph represents the percent of parent gated population for each Hardy fraction with SEM. **C)** Representative analyses for follicular and marginal zone B cells from C57BL/6 (WT) and NE1$^{-/-}$ spleens. Spleen cells were stained with antibodies (B220-PE, CD21-FITC and CD23-Biotin + Streptavidin-APC) and analyzed by flow cytometry for the presence of follicular (B220$^{+}$CD21$^{lo}$CD23$^{hi}$) and marginal zone (B220$^{+}$ CD21$^{hi}$CD23$^{lo}$) B cells. **D,E)** Graphs for follicular **(D)** and marginal zone **(E)** B cells are a representative of two independent experiments. Horizontal bar indicates the mean and each symbol represents a single mouse. P values are calculated using a two-tailed unpaired Student's t-test, ns = not significant. **F)** Representative analyses for peritoneal B1 B cells from WT and NE1$^{-/-}$ mice and stained with antibodies (CD19-PerCp, B220-APC Cy7, CD43 (S7)-APC, IgM-e450, IgD AF700, CD5-PE and vital dye FVS-510). **G)** The graphs represents at least three independent experiments. Horizontal bar indicates the mean and each symbol represents an individual mouse. P values are calculated using a two-tailed unpaired Student's t-test, ns = not significant. **H)** Sorting protocol for pro-B cell purification used in VDJ-seq. CD19$^{+}$ cells were isolated from WT and NE1-/- bone marrow using CD19-conjugated MACS beads. CD19$^{+}$ cells were stained with antibodies to CD19, CD93, CD2, CD43 and IgM. Sorted pro-B cells (CD19$^{+}$, CD93$^{+}$, IgM-, CD2$^{-}$, CD43$^{-}$) were used to isolate gDNA. Data is representative of at least three independent experiments +/- SEM.

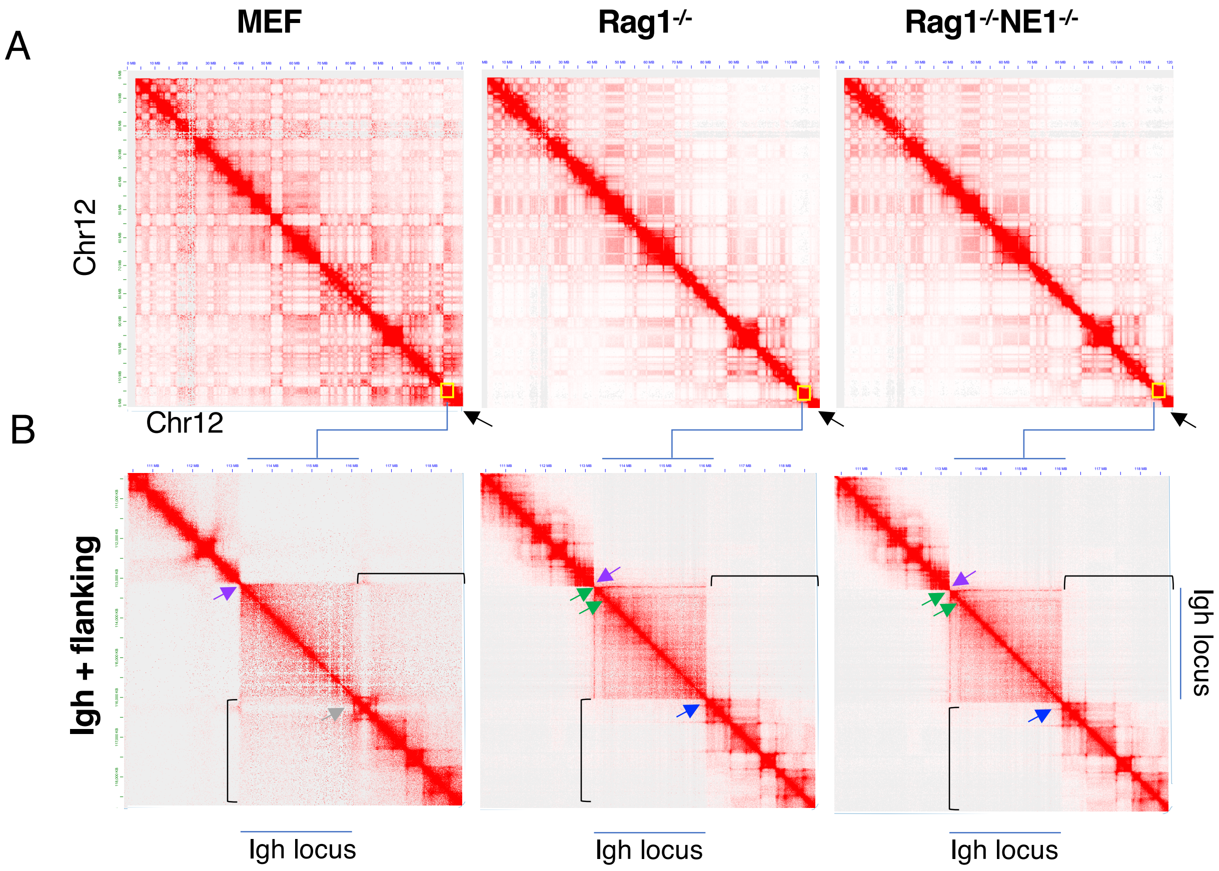


**Suppl. Figure 5. Hi-C contact maps derived from MEF, Rag1$^{-/-}$ and Rag1$^{-/-}$NE1$^{-/-}$ pro-B cells.** The Hi-C data was KR normalized. **A)** Hi-C heatmaps for chromosome 12 depicting compartments and TADs. Igh TAD (yellow box) indicated by the black arrow. **B)** Hi-C contact maps of the Igh locus and flanking regions spanning approximately chr12:110,500,000-118,500,000 (mm10). Igh locus TAD boundaries (3' end, purple arrow; 5' end, blue arrow; lost 5' boundary, gray arrow) Igh locus architectural stripes (green arrows). Compartments adjacent to the Igh locus (black brackets).

Suppl. Figure 5

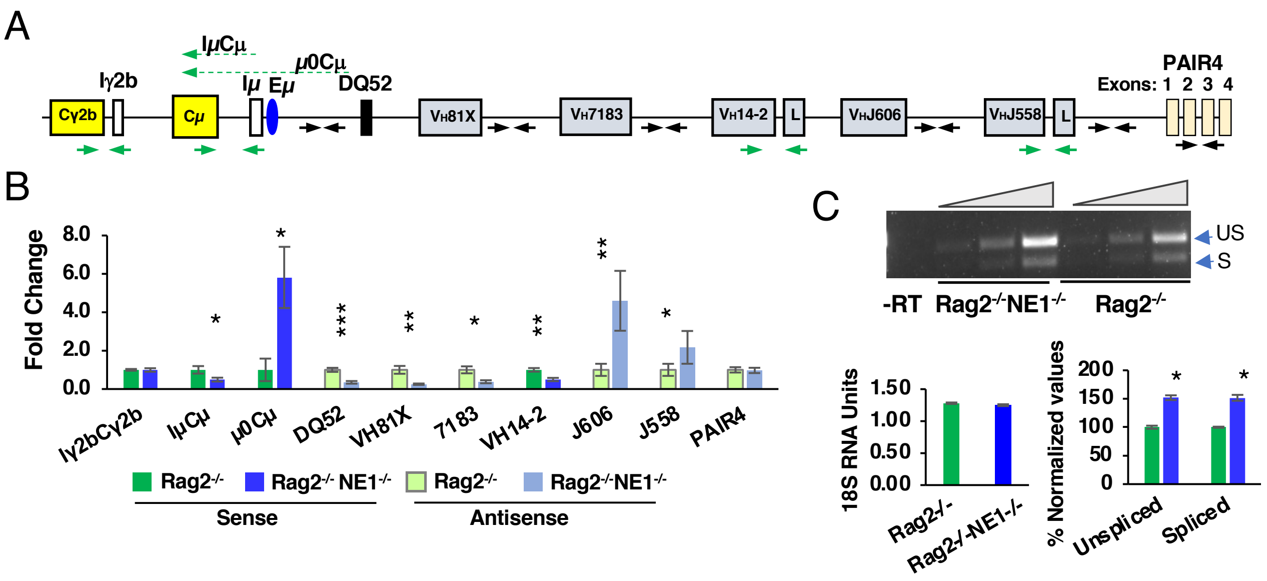


**Suppl. Figure 6. NE1 regulates $V_H$ GLT and intergenic transcription in primary Rag2$^{-/-}$- and Rag2$^{-/-}$NE1$^{-/-}$-pro-B cells. A)** Diagram for Igh locus index genes showing primers for sense (green arrows) and intergenic antisense (black arrows) transcripts. **B)** Transcripts from Rag2$^{-/-}$- and Rag2$^{-/-}$NE1$^{-/-}$- pro-B cells derived from three independent mice were analyzed in duplicate by qRT-PCR then averaged with SEMs shown. P values from Student's two tailed t test (p>0.05 (*), p>0.01 (**), p>0.001 (***)). **C)** Representative sense $V_H$J558 gene expression, that was unspliced (US, 465 bp) and spliced (S, 382 bp) was tested in semi-quantitative RT-PCR (*upper panel*) using qRT-PCR for 18S RNA as a loading control (*lower left panel*). No reverse transcription (-RT). PCR products were quantitated by densitometry and the values for Rag2$^{-/-}$ samples were normalized to 100.

**Suppl. Figure 6**
